# Bidirectional alterations in antibiotics susceptibility in *Staphylococcus aureus - Pseudomonas aeruginosa* dual-species biofilm

**DOI:** 10.1101/334516

**Authors:** Elena Y. Trizna, Maria N. Yarullina, Diana R. Baidamshina, Farida S. Akhatova, Elvira V. Rozhina, Rawil F. Fakhrullin, Alsu M. Khabibrakhmanova, Almira R. Kurbangalieva, Mikhail I. Bogachev, Airat R. Kayumov

## Abstract

While in biofilms bacteria are embedded into an extracellular matrix which forms inaccessible barrier for antimicrobials thereby drastically increasing the concentrations of antibiotics required for treatment. Here we show that the susceptibility of *S. aureus* and *P. aeruginosa* to antibiotics in mixed biofilms significantly differs from monoculture biofilms depending on both conditions and chosen antimicrobial agents. While *S. aureus* could completely avoid vancomycin, ampicillin and ceftriaxone by embedding into the biofilm of *P. aeruginosa*, the very same consortium was characterized by 10–fold increase in susceptibility to broad-spectrum antimicrobials like ciprofloxacin and aminoglycosides compared to monocultures. These data clearly indicate that efficient treatment of biofilm-associated mixed infections requires antimicrobials active against both pathogens, since the interbacterial antagonism would enhance the efficacy of treatment. Moreover, similar increase in antibiotics efficacy was observed when *P. aeruginosa* suspension was added to the mature *S. aureus* biofilm, compared to *S. aureus* monoculture, and vice versa. These findings open promising perspectives to increase the antimicrobial treatment efficacy of the wounds infected with nosocomial pathogens by the transplantation of the skin residential microflora.

## Introduction

Bacterial fouling is an important factor that strongly affects acute and chronic wounds healing and prevents wound scratch closure ^1^. Besides the physical obstruction of the cells, pathogenic bacteria produce various virulence factors including toxins and proteases that also affect cytokine production by keratinocytes, induce apoptosis of the host cells and cause inflammation ^2–6^.

*S. aureus* and *P. aeruginosa* are one of the most widespread pathogens causing various nosocomial infections, including pneumonia on the cystic fibrosis background, healthcare associated pneumonia and chronic wounds ^7–10^. During infection, bacterial cells are embedded into a self-produced extracellular matrix of organic polymers this way forming either mono- or polymicrobial biofilms ^11,12^ which drastically reduce their susceptibility to both antimicrobials and the immune system of the host ^13,14^. Current data suggests that bacterial pathogenicity is promoted during polymicrobial infections and recovery is delayed in comparison with monoculture infections ^15–17^. Accordingly, interspecies interactions between *S. aureus* and *P. aeruginosa* within mixed biofilms attracted major attention in recent years including both *in vitro* ^15^ and *in vivo* studies ^16^.

*P. aeruginosa* is known as a common dominator in polymicrobial biofilm-associated infections due to multiple mechanisms allowing its rapid adaptation to the specific conditions of the host. In particular, *P. aeruginosa* produces multiple molecules to compete with other microorganisms for space and nutrients. The main anti-staphylococcal tools of *P. aeruginosa* are siderophores and 2-*n*-heptyl-4-hydroxyquinoline *N*-oxide (HQNO), the inhibitor of the electron transport chain of *S. aureus.* Their presence shifts *S. aureus* to a fermentative mode of growth, eventually leading to reduced *S. aureus* viability ^18–22^, forces increased *S. aureus* biofilm formation ^23,24^ and transition of *S. aureus* into small-colony variants (SCVs) ^25^. SCV is a well-characterized phenotype detected in various diseases, including cystic fibrosis and device-related infections ^23–26^. SCVs appear as small, smooth colonies on a culture plate and grow significantly slower compared to wild type colonies. Remarkably, switch to the SCV phenotype improves the survival of *S. aureus* under unfavorable conditions, as it exhibits increased vancomycin and aminoglycoside resistance as well as intracellular survival ^25–29^. Prolonged co-culture with *P. aeruginosa* leads to higher proportions of stable *S. aureus* SCVs that is further increased in the presence of aminoglycosides ^25^.

*P. aeruginosa* was shown to reduce *S. aureus* during co-culture *in vitro* in both planktonic and biofilm forms^30–33^ and has been observed as the dominant pathogen in *S. aureus-P. aeruginosa* mixed infections ^16^. Nevertheless, despite of the antagonistic relationship of *S. aureus* and *P. aeruginosa* ^34,35^, several studies reported their mutual association both *in vitro* ^36,37^ and in acute and chronic wounds embedded in a mixed biofilm ^8,38–42^, with *S. aureus* typically residing on the wound surface, whereas *P. aeruginosa* being rather observed in the deep layers ^15,42–45^. Interestingly, in mixed *P. aeruginosa - S. aureus* biofilms from cystic fibrosis patients *S. aureus* was shown to be dominating during childhood, with *P. aeruginosa* prevalence increasing with aging and worsening patient prognosis ^46–48^. During the biofilm formation *P. aeruginosa* produces three main exopolysaccharides, namely alginate, Pel, and Psl, which form an extracellular matrix in the biofilm exhibiting both structural and protective functions ^49–52^. Under prevalent Pel secretion, loose biofilm structures are formed ^53^ and thus *S. aureus* is able to penetrate into the biofilm ^53^. In has been recently suggested that rare observation of *S. aureus* and *P. aeruginosa* together in diagnostic cultures of sputum of cystic fibrosis patients could be attributed to the existence of *S. aureus* as SCVs that are more difficult to detect due to their small size and fastidious growth requirements ^26,28^.

Investigations on alternative treatment options against biofilm-associated infections are largely based upon using specialized agents (such as quaternary ammonium compounds, curcumin or chlorquinaldol) or enzymatic treatment that in combinations with antibiotics provide high local drug concentrations avoiding systemic adverse effects ^54–60^. While many approaches to targeting staphylococcal biofilms were reported ^58,61–65^, only few successive ways of targeting *P. aeruginosa* are known ^60,66–68^. Among various compounds exhibiting anti-biofilm activities, the derivatives of 2(5*H*)-furanone have been reported to inhibit biofilm formation by *Staphylococci* ^69–73^. While many of these approaches exhibited promising results against staphylococcal monocultures, their efficiency against polymicrobial biofilms remains questionable. A few investigations indicated that extracellular polymeric substances forming the biofilm matrix provide protection against antibiotics to all inhabitants of the biofilm, including the non-producers, although the biofilm as a whole is weakened ^53,74^, this way proposing that *S. aureus* could potentially survive in the presence of *P. aeruginosa* and even co-exist with it in a polymicrobial biofilm, benefiting from the antimicrobial barrier formed by the *P. aeruginosa* matrix components.

Here we demonstrate that *S. aureus* successfully survives in the *P. aeruginosa* biofilm matrix under conditions of staphylococcus-specific treatment. In contrast, the efficiency of antimicrobials active against both bacterial species like ciprofloxacin and aminoglycosides in mixed biofilms increased nearly 10-fold in comparison with corresponding monocultures. These data suggest bi-directional influences on the antibiotic susceptibility in mixed biofilms, the fact that should be taken in account when considering an optimized strategy of polymicrobial infections treatment.

## Results

### Modeling the *S. aureus – P. aeruginosa* mixed biofilm

Despite of known antagonistic interactions between *S. aureus* and *P. aeruginosa* ^35^, they are still the most common pathogens evoking wound infections and forming mixed biofilms on their surfaces ^39,40,42^. We have simulated *in vitro* different situations where either *S. aureus* suspension was added to the preformed 24-h old biofilm of *P. aeruginosa* or, vice versa, *P. aeruginosa* was added to the preformed 24-h old biofilm of *S. aureus*. As a control, both strains were inoculated simultaneously and grown for 48 h with the broth exchange after 24 h of cultivation. Both *S. aureus* and *P. aeruginosa* were able to penetrate into the preformed biofilm of the other bacterium (Fig. 1). Irrespective of which bacterium initially preformed the biofilm and which one was added later, the ratio of their CFUs in the biofilm after 24 h cultivation remained around 1:10 with the prevalence of the first biofilm former (Fig. 1 A and B), and was 1:1 when both bacteria were inoculated simultaneously (Fig. 1 C).

**Figure 1.**
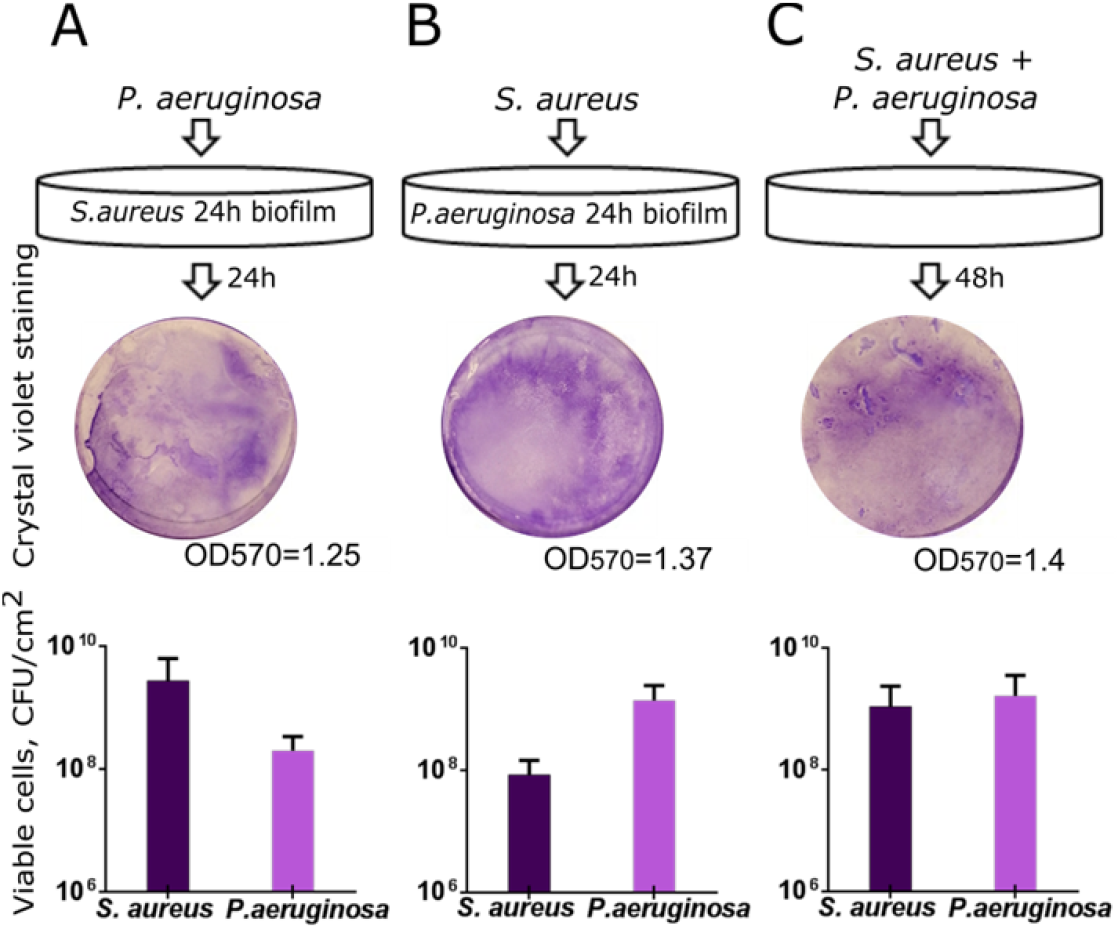
*In vitro* simulation of the *S. aureus-P. aeruginosa* mixed biofilm. (A) *P.aeruginosa* suspension in a fresh broth was added to the preformed 24-h old biofilm of *S. aureus* or (B) *S. aureus* was added to the preformed 24-h old biofilm of *P. aeruginosa* and cultivation was continued for the next 24 h. (C) As a control, both strains were inoculated simultaneously and were grown for 48 h with the broth exchange after 24 h of cultivation. The biofilms were then assessed with either crystal-violet staining of wells bottom (upper lane) or differential CFUs counting.

Next to analyze the biofilm structure and cells distribution in the matrix, the *S. aureus - P. aeruginosa* mixed biofilm was grown in imaging cover slips, live/dead stained with SYTO9/PI and analyzed with confocal laser scanning microscopy. Both *S. aureus* and *P. aeruginosa* formed 20-25 μm-thick biofilms when growing as monocultures (Fig. 2 A, B). While the mixed biofilm was of similar thickness, it appeared more rigid in comparison with monoculture ones and the fraction of non-viable cells was similar to monocultures (compare Fig. 2 A, B and C and Fig. S1 A) suggesting stability of *S. aureus - P. aeruginosa* consortium under the conditions used. By using differential staining of *S. aureus* and *P. aeruginosa* (Fig. 2 D and Fig. S1 A) we have also analyzed the distributions of *S. aureus* (red-stained) and *P. aeruginosa* (blue-stained) over the biofilm layers and evaluated their relative fractions in each layer (Fig. 2E). Interestingly, in the mixed biofilm, *S. aureus* appeared as microcolonies embedded in the biofilm matrix (see white arrow in Fig. 2 C and 2 D) tended to distribute in the upper layers of the biofilm, while *P. aeruginosa* dominated in its lower layers (see Fig. 2 E).

**Figure 2.**
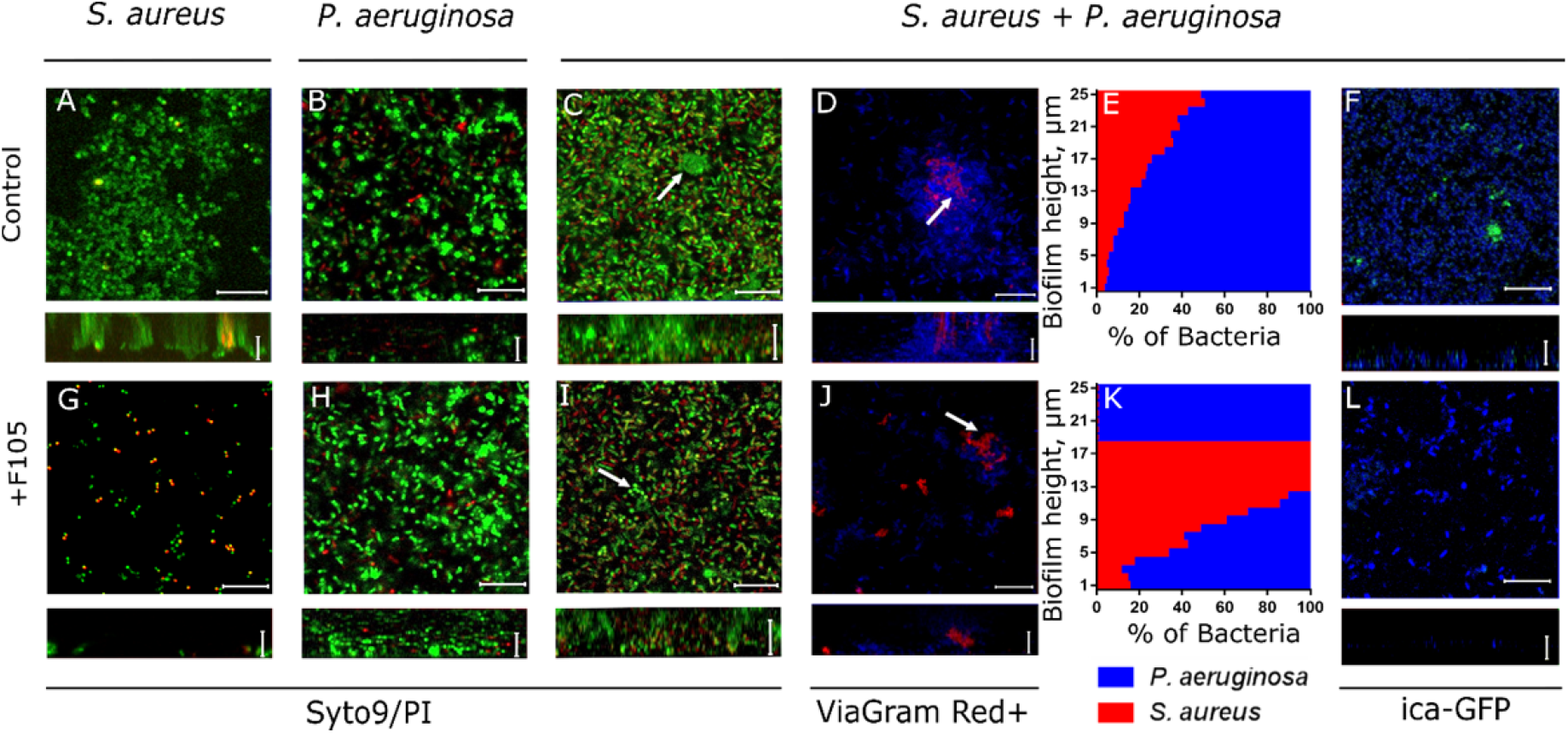
Mono- and polymicrobial biofilms formed by *S. aureus* and *P. aeruginosa*. Cells were grown without any antimicrobial (A-F) or in presence of 2(5*H*)-furanone derivative (F105, G-L) specifically inhibiting the biofilm formation by *S. aureus* and exhibiting no effects on *P. aeruginosa*. The 48-h old biofilms were stained by Syto9/PI (A-C, G-I) or ViaGram Red^+^ (D, J) to differentiate *S. aureus* (stained in red) and *P. aeruginosa* (stained in blue) and assessed by CLSM. (E, K) The distribution of *S. aureus* and *P. aeruginosa* in the biofilm layers expressed as their relative fractions. The repression of *S. aureus ica*-operon in mixed biofilm by F105 was monitored by detection of GFP in *ica*-GFP strain (F, L). The scale bars indicate 10 μm. *S. aureus* microcolonies in a mixed biofilm are shown by arrows.

In the last decades different approaches to inhibit the biofilm formation by various bacteria were developed ^55,56,58^, appearing nowadays more successful in prevention of *S. aureus* biofilm formation ^61,62,64^. Therefore we simulated the *S. aureus - P. aeruginosa* mixed biofilm formation under the conditions of biofilm-preventing treatment. For that, bacteria were cultivated in the presence of a derivative of 2(5*H*)-furanone denoted as F105, identified in recent study as an efficient inhibitor of biofilm formation by *S. aureus* ^73,75^, while exhibiting no significant effect against *P. aeruginosa* (Table S1). When *S. aureus* was grown for 48 h in the presence of 2.5 μg/ml of F105, no biofilm was formed and cells growth was retarded, while most of the cells remained viable in both surface-adherent (Fig. 2 G) and planktonic state (Fig. S2 A). As well, the repression of the biofilm formation was confirmed by evaluation of the ica-GFP expression in presence of F105. No GFP was observed in cells grown in presence of F105 in *S.aureus* pRB-ica-gfp (Fig. S3 C, D), while the constitutive expression of GFP in *S.aureus* pC-tuf-gfp^76^ was not affected under these conditions (Fig. S4 B, D) confirming the inhibition of the biofilm formation. As expected, no significant effect of F105 on *P. aeruginosa* viability could be observed (Fig. 2 H, Table S1). Moreover, the biofilm formation was slightly increased as determined by crystal violet staining (Fig. S5) and CLSM (compare Fig. 2 B and H).

When *S. aureus* and *P. aeruginosa* were grown together in the presence of F105, *S. aureus* microcolonies were also observed, similarly to the control (compare Fig. 2 C, D and I, J), suggesting that *S. aureus* cells are able to form clusters inside the biofilm of *P. aeruginosa*, despite of its antagonistic pressure (see white arrows on Fig. 2 I, J). No GFP in *S. aureus* pRB-ica-gfp was observed (Fig. 2 L), indicating that in presence of F105 in mixed biofilm the matrix is produced mainly by *P. aeruginosa*. Next, in marked contrast to the control, the *S. aureus* cells were observed predominantly in the middle layers of the biofilm (compare Fig. 2 E and K).

The microscopic data were also validated by direct CFU counting in the biofilm; by using mannitol salt agar plates and cetrimide agar plates the bacterial species were differentiated and their CFUs were counted separately (Fig. S2). In the presence of F105 the amount of adherent viable *S. aureus* cells decreased by six orders of magnitude in monoculture, suggesting complete inhibition of the biofilm formation, while no significant differences in CFUs of *P. aeruginosa* could be observed. In a mixed biofilm, the *S. aureus* to *P. aeruginosa* ratio remained unchanged in the control, while the fraction of viable *S. aureus* cells decreased slightly in the presence of F105, this way confirming CLSM data and supporting the hypothesis that *S. aureus* is apparently able to embed and hide thereby in the biofilm formed by *P. aeruginosa* when its own biofilm formation is repressed.

### Atomic force microscopy

The atomic force microscopy of both monocultures and mixed biofilms of *S. aureus - P. aeruginosa* confirmed the CLSM data. Thus, in control wells the biofilms of monocultures of both strains formed a typical confluent multilayer biofilm (Fig. 3 A, B), in mixed biofilm *S. aureus* was prevalently distributed in the upper layers (Fig. 3 C). Interestingly, the adhesion force of the mixed biofilm was 3-fold lower compared to *S. aureus* monoculture biofilm and 2-fold lower compared to *P. aeruginosa* monoculture biofilm (Fig. 4), suggesting more irregular structure of the mixed biofilm ^53^. When growing with F105, only *P. aeruginosa* could be observed on the biofilm surface in the mixed culture, suggesting that *S. aureus* was hidden into the lower biofilm layers. Since the adhesion force of the mixed biofilm in the presence of F105 was similar to that one in the monoculture *P. aeruginosa* (Fig. 3 F and Fig. 4), we assumed that the biofilm matrix under these conditions was presumably formed by *P. aeruginosa*.

**Figure 3.**
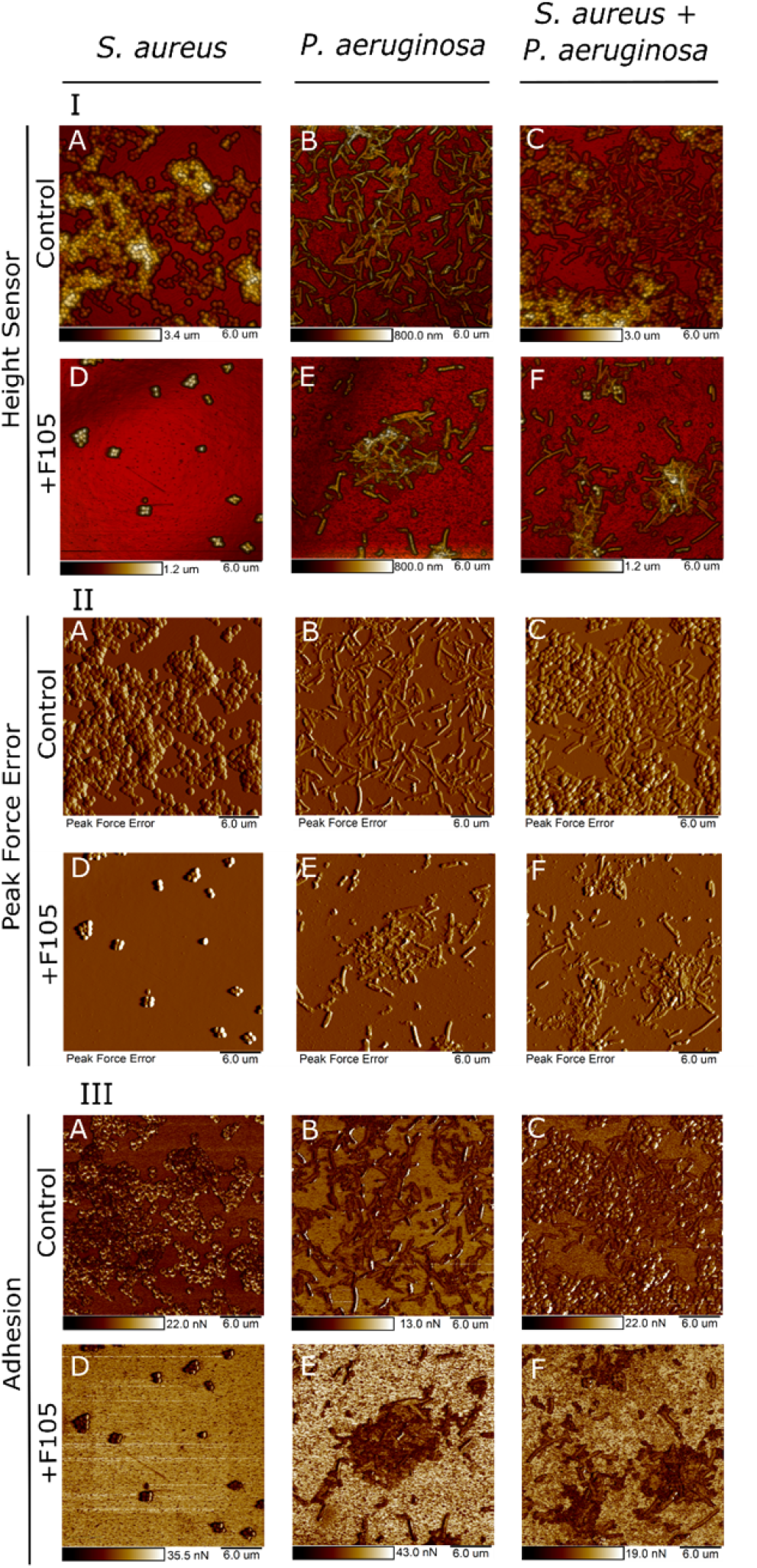
Atomic force microscopy (Peak Force Tapping mode) of mono- and polymicrobial biofilms formed by *S. aureus* and *P. aeruginosa*. Cells were grown without any antimicrobials (A-C) or in presence of F105 specifically inhibiting the biofilm formation by *S. aureus* cells (D-F) for 48 hours, then the plates were washed, fixed with glutardialdehyde and analyzed with AFM. (I) – Sensor height (topography); (II) – 3D reconstruction of height channel image; (III) –adhesion.

**Figure 4.**
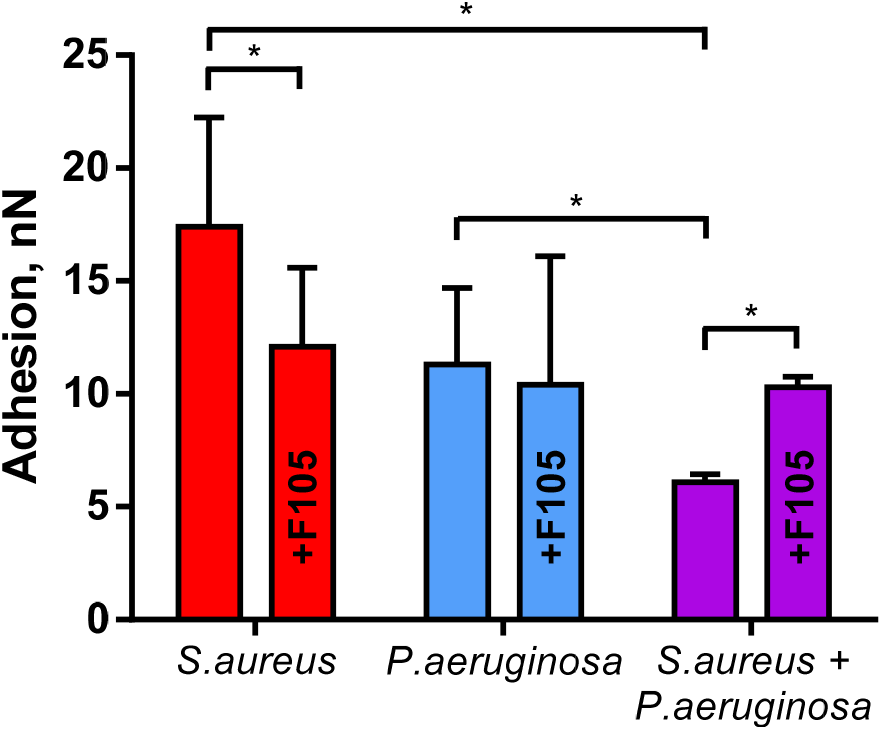
The adhesion force of *S. aureus* and *P. aeruginosa* monoculture and mixed biofilms. To repress the biofilm formation F105 was added up to 2.5 μg/ml. Asterisks denote statistical significant difference was confirmed by the Kruskal-Wallis statistical test at *p* < 0.05.

### *S. aureus* and *P. aeruginosa* susceptibility to antibiotics in mixed biofilms

Our data suggest that *S. aureus* under anti-biofilm treatment conditions is able to form microcolonies in the biofilm of *P. aeruginosa*, thereby apparently changing their susceptibility to antimicrobials. To further verify this assumption, the effect of various conventional antibiotics on preformed mono- and polymicrobial biofilms was studied. The 48-h old monoculture and mixed biofilms were prepared in 24-well adhesive plates in either absence or presence of F105 to repress the biofilm formation by *S. aureus* itself. Then the biofilms were washed with sterile 0.9% NaCl and wells were loaded with fresh broth supplemented with antibiotics at wide range of final concentrations to fill the range of their 1-16 fold MBCs (see table S1 for MBC values). After 24h incubation the amount of CFUs of both *S. aureus* and *P. aeruginosa* in both the culture liquid and the biofilm was determined by the drop plate assay and the distribution of cells in the mixed biofilm was assessed by CLSM.

First, the biofilm-eradicating activity was investigated for the antibiotics conventionally used for *S. aureus* treatment but typically inefficient against *P. aeruginosa* including vancomycin, tetracycline, ampicillin and ceftriaxone (Figs 5, S6, S7). In monoculture, vancomycin reduced the amount of viable *S. aureus* cells in the biofilm by 3 orders of magnitude only at 16×MBC (Fig. 5 A). In the culture liquid, 0.125×MBC (corresponds to 1×MIC, see Table S1) of vancomycin was sufficient to decrease of CFUs count by three orders of magnitude (Fig S6). Apparently, it could be attributed to the protection of detached cell clumps with EPS tailings (Fig. S8). Expectedly, in the presence of F105 (2.5 μg/ml) no biofilm could be formed and bacteria were found completely dead both in the biofilm and in culture liquid after 24-h exposition to the antibiotic at 1×MBC (Fig. 5 C, S6). Irrespective of either presence or absence of F105 *P. aeruginosa* remained resistant to the antibiotic (Fig. 5 B, D).

**Figure 5.**
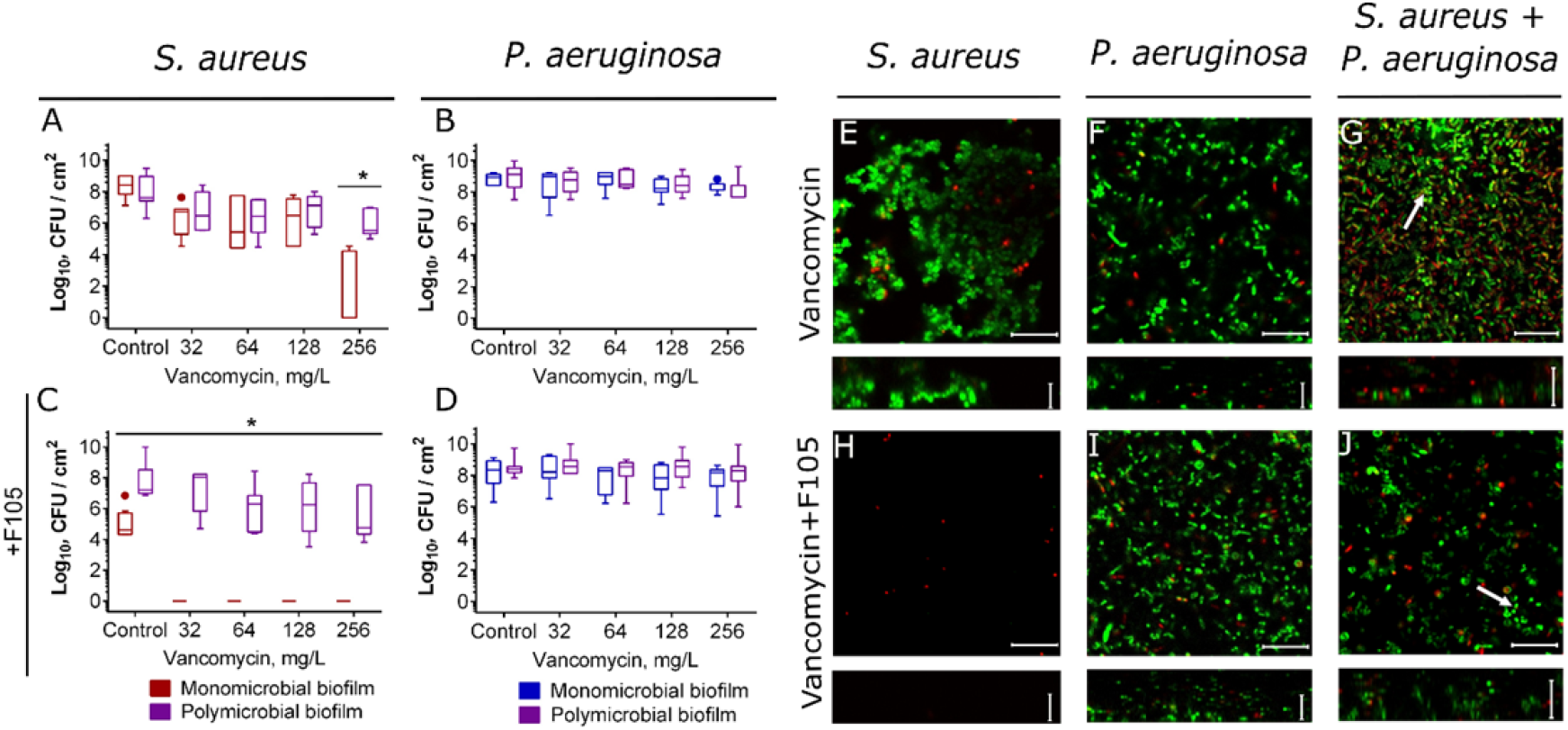
The effect of vancomycin on viability of *S. aureus* and *P.aeruginosa* embedded into their mono- and polymicrobial biofilms. Vancomycin was added to 48 hours-old biofilms grown in absence (A-B, E-F) or presence (C-D, H-J) of F105 to inhibit the biofilm formation by *S. aureus*. After 24 h incubation, the biofilms were washed twice with sterile 0.9% NaCl. The adherent cells were scratched, resuspended and CFUs were counted. Alternatively, biofilms were stained by Syto9/PI and biofilms were assessed by CLSM. The scale bars indicate 10 μm.

In a mixed culture, irrespective of the *S. aureus* biofilm formation repression by F105, viable *S. aureus* cells were identified within the biofilm and the efficiency of antibiotic reduced drastically (Figs. 5 A, C, compare reds and violets). Statistical significance of this discrepancy was confirmed by the Kruskal-Wallis statistical test at *p* < 0.05. Similar inefficiency of vancomycin against *S. aureus* in the culture liquid was observed in mixed cultures (Fig. S6).

For a deeper understanding of localization distribution and viability of bacteria in mixed biofilms under vancomycin treatment, the CLSM analysis has been performed. In the presence of F105 no biofilm of *S. aureus* could be observed resulting in significant decrease of viable cells fraction after vancomycin treatment, in contrast to the biofilm-embedded cells (compare Fig. 5 E and H). In the mixed biofilm the cell clusters (apparently *S. aureus* microcolonies) were observed in the biofilm matrix similarly to the control (compare Fig. 2 C, I and Fig 5 G, J), suggesting that *staphylococci* are able to escape the antimicrobials and survive by embedding into the polymicrobial biofilm. In marked contrast to the control where *S. aureus* was mostly located in the top layers of the biofilm, under vancomycin treatment most of *S. aureus* cells appeared in the lower and middle layers of the biofilm (compare Fig. 2 E and Fig. 6). These data suggest that vancomycin-resistant *P. aeruginosa* cells in the upper layers of the biofilm apparently prevented the penetration of the antibiotic into the matrix this way reducing the susceptibility of *S. aureus* to antibiotics considerably. Of note, *S. aureus* cells remained mainly viable in bottom layers apparently because of protection by *P. aeruginosa* cells (Fig. S9).

**Figure 6.**
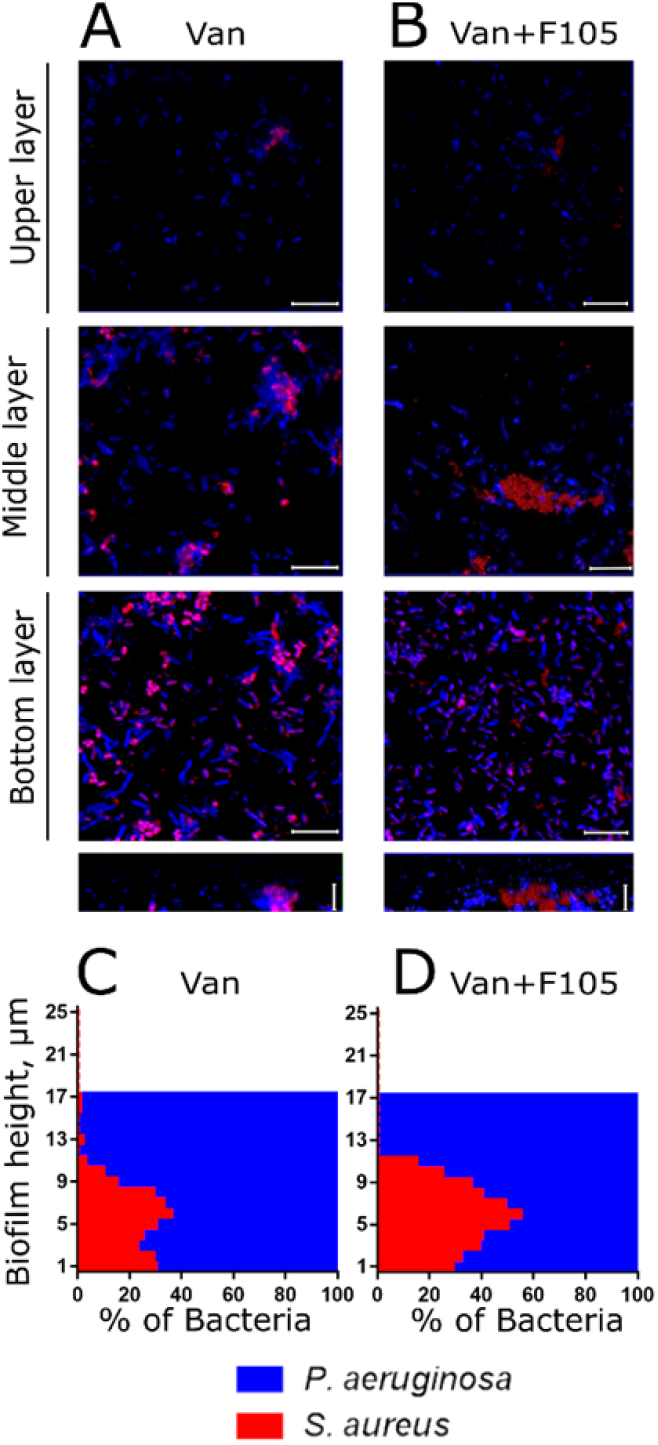
The effect of vancomycin on *S. aureus* and *P. aeruginosa* distribution in mixed biofilms. Cells were grown in absence (A, C) or in presence of F105 specifically inhibiting the biofilm formation by *S.aureus* cells (B, D). Vancomycin (256 μg/mL corresponding to 8×MBC for *S. aureus*) was added to 48 hours-old biofilms. After 24 h incubation, the biofilms were stained by ViaGram Red^+^ to differentiate *S. aureus* (stained in red) and *P. aeruginosa* (stained in blue) and assessed by CLSM. The scale bars indicate 10 μm. (C, D) The distribution of *S. aureus* and *P. aeruginosa* in the biofilm layers are expressed as their relative fractions.

It has been suggested previously that 2-n-heptyl-4-hydroxyquinoline N-oxide (HQNO) and siderophores pyoverdine and pyochelin produced by *P. aeruginosa* force the shift of *S. aureus* to a fermentative mode of growth under anoxia and decrease thereby their susceptibility to vancomycin, the cell wall-targeting antibiotic^29^. To test whether this mechanism or hiding into the biofilm of *P. aeruginosa* primarily govern the decreased susceptibility of *S. aureus* to antimicrobials in the mixed biofilms, the effect of other conventional antibiotics effective only against *S. aureus* was investigated. Thus, ampicillin and ceftriaxone, antibiotics also targeting the cell wall synthesis although by another mechanism (by irreversible inhibition of the transpeptidase), as well as tetracycline, which inhibits the translation initiation by binding to the 30S ribosomal subunit, were used.

Similarly to vancomycin, treatment by ampicillin, tetracycline and ceftriaxone was almost inefficient against biofilm-embedded *S. aureus*, while under conditions of biofilm formation repression by F105, the 1-2×MBC of antimicrobials led to the complete death of cells in 24 h (Fig. S7). In culture liquids, 0.25-0.5×MBC of ampicillin and ceftriaxone provided the full death of *S. aureus*, 0.125×MBC of tetracycline decreased the amount of viable *S. aureus* cells by 3 orders of magnitude. F105 reduced by factor 4 the amount of antimicrobials sufficient to kill bacteria completely (Fig. S7 A, C). In the mixed culture, even despite of the *S. aureus* biofilm formation repression with F105, in both the culture liquid and biofilm *S. aureus* cells remained insensitive to any of antimicrobials tested (Fig. S7 A, C). CLSM analysis indicated considerable redistribution of *S. aureus* from upper to the bottom layers of the biofilm (Fig. S10), apparently leading to reduced antibiotic efficacy. Under double treatment by F105 and antimicrobials, the prevalence of *P. aeruginosa* in the biofilm was observed in agreement with the CFU count data (Figs S7 and S10).

These data suggest that under antimicrobial treatment conditions *S. aureus* changes its preferred topical localizations by hiding in the lower layers of mixed biofilm formed by another bacterium like *P. aeruginosa* insensitive to most antimicrobials probably thereby increasing its resistance to the treatment.

Next, we investigated the effect of broad-spectrum antimicrobials such as ciprofloxacin, amikacin and gentamycin which are active against both *S. aureus* and *P. aeruginosa* (see Table S1). In contrast to the previous group of antimicrobials, high concentrations of ciprofloxacin efficiently eradicated both *P. aeruginosa* and *S. aureus* monocultures even in the biofilm-embedded form (Fig. 7 A, B). Interestingly, when the mixed biofilm was treated, nearly 10-fold lower concentration of antimicrobial was sufficient to obtain similar reduction of *P. aeruginosa* CFUs in the biofilm, although not for detached cells (Fig 7 B, Fig S11). Moreover, significant increase of susceptibility of both biofilm-embedded and detached *S.aureus* to ciprofloxacin was observed in mixed culture in contrast to monoculture (see Figs 7A, S11 A). Finally, in the mixed biofilm the complete death of both *P. aeruginosa* and *S. aureus* could be observed at 8×MBC of ciprofloxacin, in marked contrast to monocultures (Fig 7 A, B).

**Figure 7.**
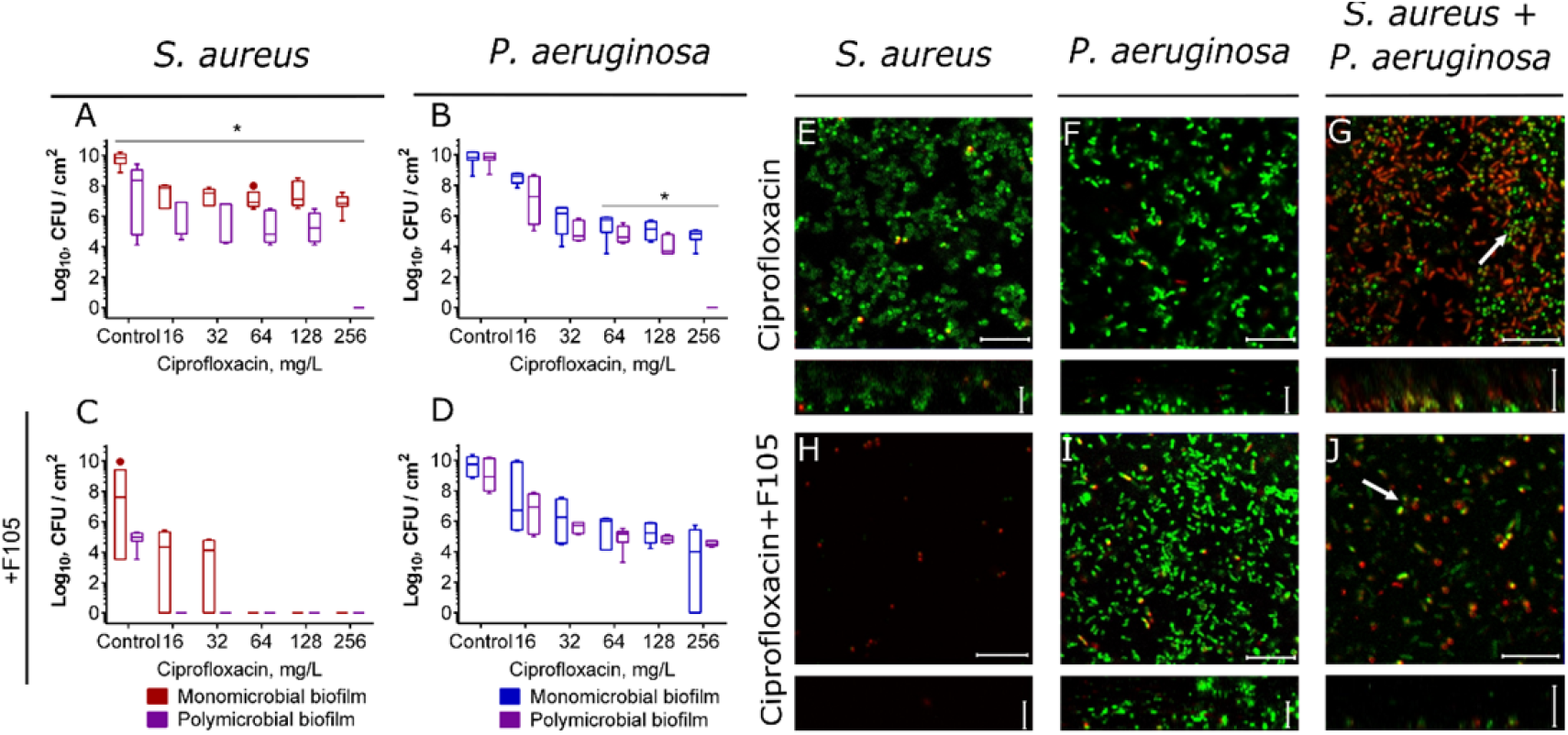
The effect of ciprofloxacin on viability of *S. aureus* and *P.aeruginosa* embedded into their mono- and polymicrobial biofilms. Ciprofloxacin was added to 48 hours-old biofilms grown in absence (A-B, E-F) or presence (C-D, H-J) of F105 to inhibit the biofilm formation by *S. aureus*. After 24 h incubation, the biofilms were washed twice with sterile 0.9% NaCl. The adherent cells were scratched, resuspended and CFUs were counted. Alternatively, biofilms were stained by Syto9/PI and biofilms were assessed by CLSM. The scale bars indicate 10 μm.

Similarly, 1-2×MBC of aminoglycosides (amikacin or gentamicin) led to the complete death of both *P. aeruginosa* and *S. aureus* in mixed biofilm while reducing their CFUs in monocultures only by 2-3 orders of magnitude at 8×MBCs (Fig. 8). For detached *S. aureus* cells the effect was less pronounced (Fig. S12), while the susceptibility of *P. aeruginosa* to both aminoglycosides in the culture liquid was slightly increased in contrast to ciprofloxacin (Fig. S11).

**Figure 8.**
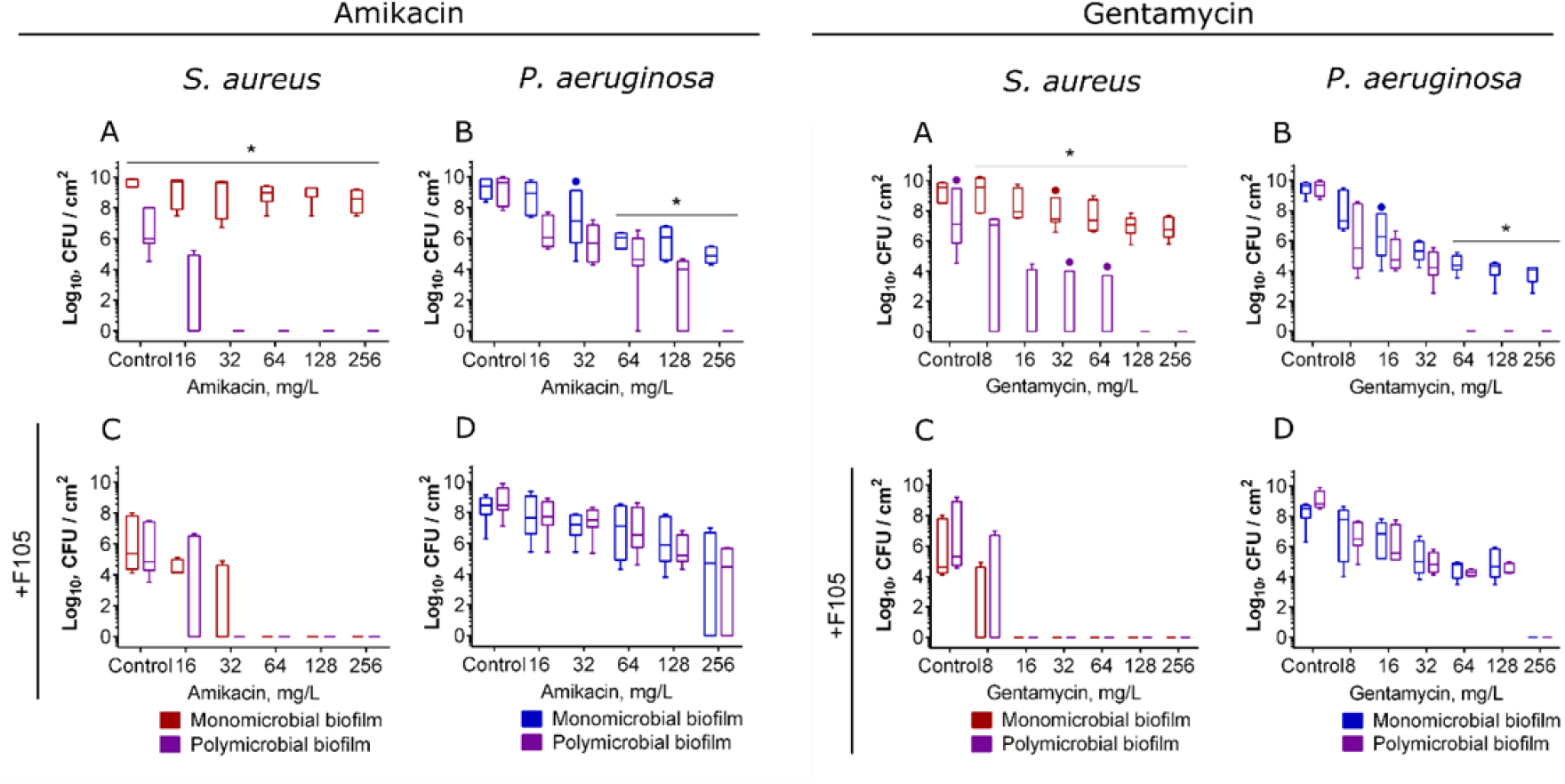
The effect of amikacin and gentamycin on viability of *S. aureus* and *P.aeruginosa* in mono- and polymicrobial biofilms. Antimicrobials were added to 48 hours-old biofilms grown in absence (A-B) or presence (C-D) of F105 to inhibit the biofilm formation by *S. aureus*. After 24 h incubation, the biofilms were washed twice with sterile 0.9% NaCl. The adherent cells were scratched, resuspended and CFUs were counted.

In the presence of F105, just 0.5×MBC of any tested antimicrobial was already sufficient for the complete death of both *S. aureus* detached and biofilm-embedded cells (see Fig. 5 C and Fig. 7 C), similarly to vancomycin, tetracycline, ampicillin and ceftriaxone (see Fig. 5 and Fig. S7). The presence of F105 did not affect the susceptibility of monoculture *P. aeruginosa* biofilm to antibiotics. In contrast, in mixed cultures the inhibition of *S. aureus* by F105 restored the susceptibility of *P. aeruginosa* back to the monoculture level, suppressing the observed high efficiency of antimicrobials against this bacterium in the mixed biofilm (compare pallets B and D in Figs. 7, 8, S11, S12). Surprisingly, the efficiency of aminoglycosides against *S. aureus* in mixed culture did not depend on presence of F105 (pallets C in Figs. 7, 8, S11, S12).

The CLSM analysis of *S. aureus* and *P. aeruginosa* monoculture and mixed biofilms treated with ciprofloxacin confirmed the CFUs counting data. In particular, while 8×MBC did not affect either *S. aureus* or *P. aeruginosa* cells in monoculture biofilms (Fig. 7 E, F), in the mixed biofilm a huge fraction of non-viable cells was observed (Fig. 7 G). In marked contrast, repression of the *S. aureus* biofilm production by F105 led to a reversal with most *P. aeruginosa* cells green-stained while *S. aureus* identified as non-viable in mixed culture (Fig. 7 J).

The distribution of bacteria in the mixed biofilm layers under treatment with ciprofloxacin was also assessed (Fig. 9). In contrast to vancomycin treatment, here *S. aureus* dominated in the upper layers of the mixed biofilm (compare Fig 6 and 9 A and C) and remained alive, while *P. aeruginosa* were presumably dead (see Figs S9, S13, S14) suggesting no reversal protection of *P. aeruginosa* by *S. aureus* biofilm. On the other hand, double treatment by ciprofloxacin combined with F105 resulted in hiding of *S. aureus* in the bottom layers of the biofilm and increased resistance of *P. aeruginosa*. Treatment by amikacin and gentamycin led to considerably different distributions of bacteria over the biofilm layers with the prevalence of *S. aureus* in the bottom layers irrespective of its biofilm repression by F105 (Fig. S14, cells distribution patterns) but qualitatively similar bacterial survival patterns (see Fig. S12). Moreover, under single antibiotic treatment *P. aeruginosa* were presumably dead, while *S. aureus* remained viable (Fig. S14). In the presence of F105 *P. aeruginosa* remained alive and much less *S. aureus* cells could be observed in the biofilm, as almost all of them were identified as non-viable.

**Figure 9.**
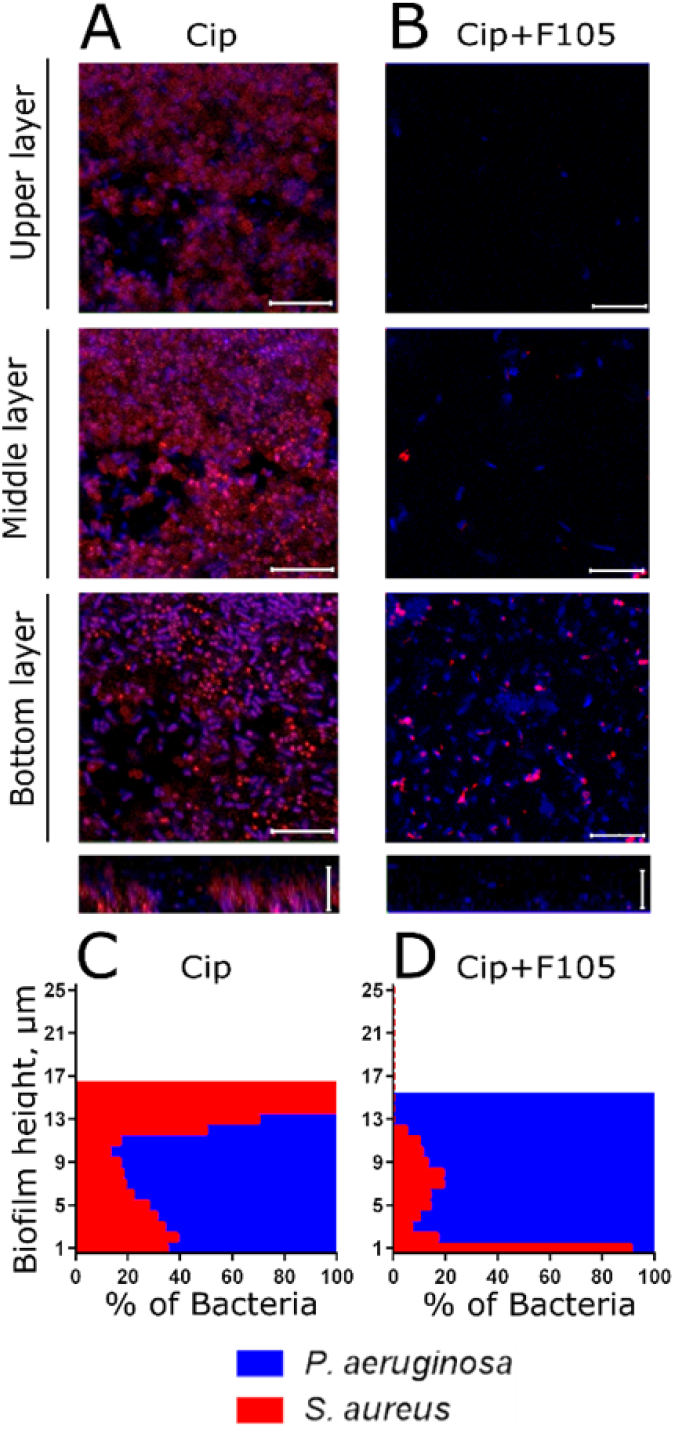
The effect of ciprofloxacin on *S. aureus* and *P. aeruginosa* distribution in mixed biofilms. Cells were grown normally (A, C) or in presence of F105 specifically inhibiting the biofilm formation by *S. aureus* cells (B, D). Ciprofloxacin (512 μg/mL corresponding to 8×MBC for both bacteria) was added to 48 hours-old biofilms. After 24 h incubation, the biofilms were stained by ViaGram Red^+^ to differentiate *S. aureus* (stained in red) and *P. aeruginosa* (stained in blue) and assessed by CLSM. The scale bars indicate 10 μm. (C, D) The distribution of *S. aureus* and *P. aeruginosa* in the biofilm layers are expressed as their relative fractions.

Taken together these data suggest that complex interspecies interactions between *S. aureus* and *P. aeruginosa* in mixed biofilm significantly affect the efficacy of treatment by antimicrobials with different specificity.

Recent data indicate that *S. aureus* forms so-called Small Colony Variants (SCV) when growing with *P. aeruginosa* or when treated with aminoglycosides due to the defects in respiration ^23–26,29^. Cells with SCV-phenotype are much more robust against external stresses compared to regular *S. aureus* cells, and thus the observed drastic decrease of *S. aureus* susceptibility to antimicrobials in mixed biofilms could be potentially attributed to the SCV formation. To verify this hypothesis, next the CFUs were counted by plating the cells on LB broth with colistin, which specifically kills *P. aeruginosa* but does not affect *S. aureus*. While the SCVs constituted up to 90% of *S. aureus* population (Fig. S15), that fits with previous reports^27^, the effect of higher efficiency of broad-spectrum antibiotics against mixed biofilms could nevertheless be observed (Fig. S15).

This fact together with the observation that the repression of *S. aureus* by F105 abrogated the effect of higher efficiency of aminoglycosides against *P. aeruginosa* in mixed biofilms indicates that the increase of broad-spectrum antibiotics efficacy against *P. aeruginosa* and *S. aureus* in mixed biofilms is less related or unrelated to SCV formation and rather governed by interspecies interaction between these bacteria. One of the main antagonistic tools of *P. aeruginosa* is the production of cyanide ^77^. To test whether it could be one of mechanisms increasing the efficiency of antimicrobials in mixed cultures, the experiments have been performed by using *S.aureus* 0349 pCXcydAB_sa_ ^77^ strain insensitive to cyanide. Despite of cyanide resistance, similar drastic reduction of *S. aureus* CFU in mixed biofilms could be observed when treated with ciprofloxacin and aminoglycosides also ruling out the cyanide synthesis by *P. aeruginosa* as the key contributing mechanism and indicating that other factors lead the observed effects (Fig. S16).

### Intervention of *P. aeruginosa* into *S. aureus* biofilm and vice versa as a possible way to enhance antimicrobial susceptibility

Our results indicate that under appropriate conditions both *S. aureus* and *P. aeruginosa* due to their antagonistic interactions appear more susceptible to broad-spectrum antimicrobials in polymicrobial biofilms, compared to their monoculture counterparts. Based on these data, we have suggested that also the susceptibility of monoculture biofilms could be increased by deliberate intervention of *P. aeruginosa* into preformed *S. aureus* biofilm, and vice versa.

To verify the efficacy of this approach, *P. aeruginosa* suspension (10^6^ CFU/mL) was added to the 24 h-old *S. aureus* biofilm and bacteria were incubated for the next 24 h. Then the biofilm was washed by sterile saline and fresh broth containing different antimicrobials was added into the wells. After 24 h the number of *P. aeruginosa* and *S. aureus* CFUs was counted by using differential media.

The introduction of *P. aeruginosa* into *S. aureus* biofilm did not change the efficacy of any antibiotic against *P. aeruginosa* itself (Fig. 10, lane II). In contrast, 1×MBC of ciprofloxacin led to the reduction of viable *S. aureus* in biofilm by 3 orders of magnitude, while in the monoculture 4-8×MBC was required to achieve the same effect (Fig. 10, lane I, compare reds and violets). Amikacin and gentamycin, being almost inefficient against *S. aureus* monoculture biofilm up to 8×MBC, were able to decrease the *S. aureus* CFUs in biofilm by 3 orders of magnitude already at 1-2×MBC after introduction of *P. aeruginosa* with the most pronounced effect observed for gentamycin.

**Figure 10.**
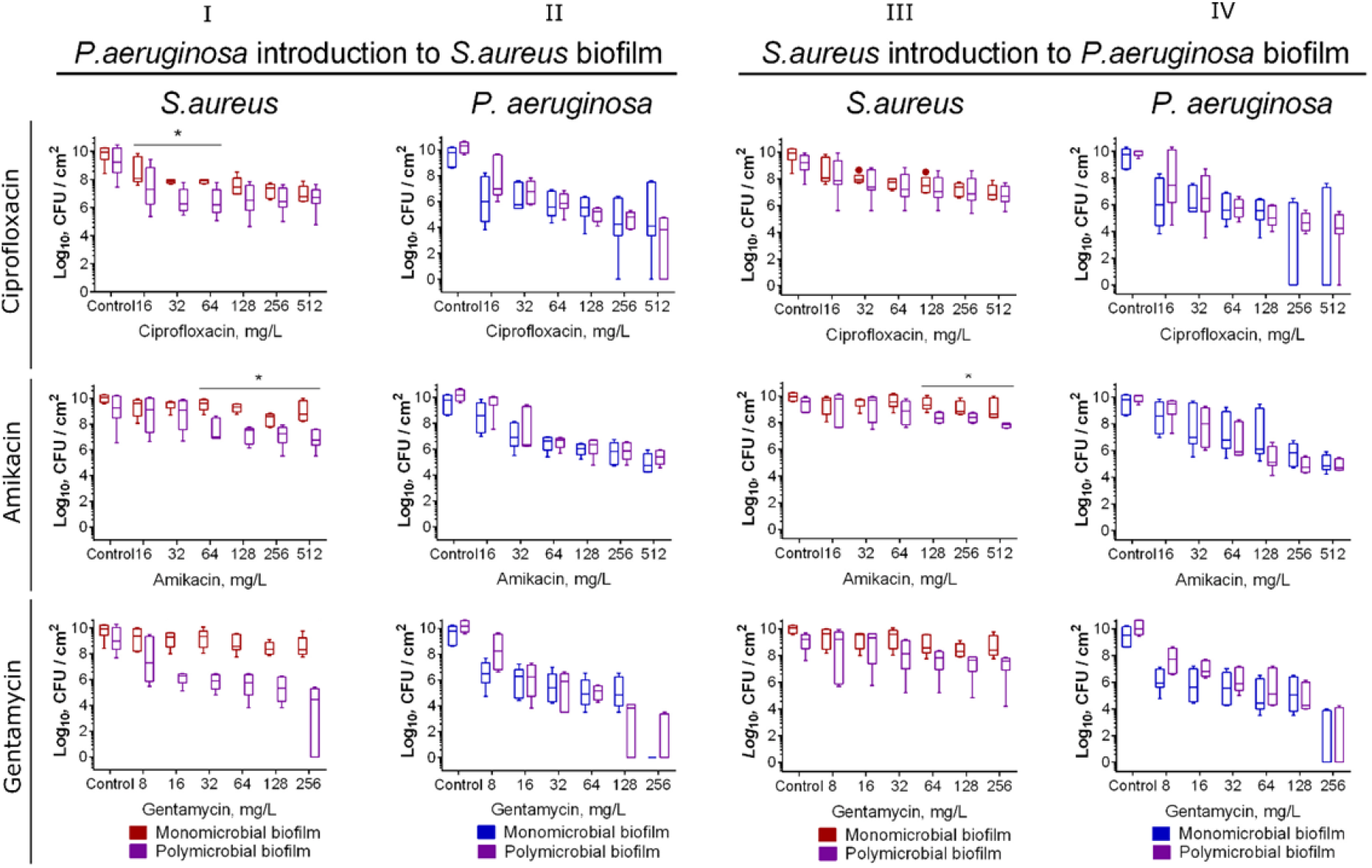
The susceptibility of *P. aeruginosa* and *S. aureus* after introduction of the antagonist into monoculture biofilms. (I-II) *P. aeruginosa* suspension in a fresh broth was added to the preformed 24-h old biofilm of *S. aureus* or (III-IV) *S. aureus* was added to the preformed 24-h old biofilm of *P. aeruginosa* and cultivation was continued for the next 24 h. Then antimicrobials were added and after 24 h incubation CFUs in biofilms were counted. Asterisks show significant difference between CFUs number between monoculture and mixed biofilms.

In the reverse experiment, when *S. aureus* was added to the *P. aeruginosa* biofilm, a remarkable increase of ciprofloxacin efficacy against *P. aeruginosa* could be observed (Fig. 10, lane IV, compare blues and violets), while the susceptibility of *S. aureus* itself did not change significantly. The efficacy of aminoglycosides has increased only against *S. aureus*, while not against *P. aeruginosa*.

## Discussion

Biofilm formation represents an important virulence factor of many bacteria, as the extracellular matrix drastically reduces their susceptibility to antimicrobials resulting in up to 1000-fold higher tolerance to antibiotics of biofilm-embedded cells compared to their planktonic forms ^14,78,79^. In contrast, polymicrobial communities are often characterized by concurrent interspecies interactions that likely overwhelm the potential benefits from biofilm protection. Here we show that in *S. aureus* – *P. aeruginosa* mixed biofilms, the most common pathogenic agents causing various nosocomial infections ^7–9^, bacterial susceptibility to antibiotics is drastically changed making them significantly more or less vulnerable to treatment than in monoculture biofilms depending on both conditions and chosen antimicrobial agents.

Despite of the antagonistic relationship between *S. aureus* and *P. aeruginosa* described in multiple studies ^34,35,80^, these bacteria can be found in close association in acute and chronic wounds being embedded into mixed biofilms ^8,38–42^. Our *in vitro* data show that the inoculation of *S. aureus* to the mature *P. aeruginosa* biofilm or vice versa leads to the formation of mixed biofilm, although with the prevalence of the first biofilm former (Fig. 1). The co-cultivation of both bacteria results in the formation of a more rigid biofilm, where *S. aureus* is located mainly in the upper layers, while *P. aeruginosa* can be found mostly in the lower layers of the biofilm (Fig. 2), in agreement with earlier data ^15,42–45^.

In several works the alteration of *S. aureus* susceptibility to various antimicrobials in presence of *P. aeruginosa* has been shown mainly in liquid cultures ^21,81,82^. Here, we investigated the effect of two groups of antimicrobials on bacterial viability in mixed biofilms. The first group contained vancomycin, tetracycline, ampicillin and ceftriaxone that are known to exhibit specific activity against *S. aureus* while leaving *P. aeruginosa* nearly unaffected. The second group included broad-spectrum antibiotics such as ciprofloxacin, gentamicin and amikacin that exhibited comparable MBC values against both studied bacteria (see table S1). Additionally, we also simulated the biofilm-preventing treatment with earlier described compound F105, specifically affecting only *S. aureus* biofilm formation ^73^. In control experiments with *S. aureus* monoculture biofilms, none of the antimicrobials exhibited any bactericidal effect at their 8-16×MBCs, while 1×MBC was already sufficient for the complete eradication of both adherent and detached cells under biofilm repression conditions with F105 (compare Figs 5, 7, 8, S6, S7, S11, S12 reds on panels A and C). In addition, ciprofloxacin, gentamicin and amikacin at 8×MBCs significantly reduced the number of CFUs of biofilm-embedded *P. aeruginosa* (Figs 5, 7, 8, S6, S7, S11, S12 blues on panels B and D).

In mixed biofilms, *S. aureus* became significantly less susceptible to antimicrobials active specifically against *S. aureus* such as vancomycin, tetracycline, ampicillin and ceftriaxone independently of presence of the biofilm repressing agent F105. It has been reported previously that HQNO produced by *P. aeruginosa* decrease the *S. aureus* susceptibility to vancomycin in mixed cultures in concentration-dependent manner and this effect was attributed to the fermentative mode of ETC action ^29^. In our experiments, not only vancomycin but the other *S. aureus-*specific antimicrobials like ampicillin, tetracycline and ceftriaxone became inefficient against *S. aureus* in dual-species biofilm (see Figs 5, S7) irrespective of staphylococcal biofilm repression with F105 (Fig. 5). CLSM analysis revealed that *S. aureus* was re-localized under treatment conditions to the middle and lower layers of the biofilm and remained viable apparently being embedded into the matrix of *P. aeruginosa* biofilm (see Figs 6, S9, S10). These data allowed suggesting that *staphylococci* are able to escape the antimicrobials by embedding into the biofilm matrix of *P. aeruginosa* and survive there, despite of antagonistic interactions between these bacteria. Notably, in mixed biofilms *S. aureus* formed cell clumps in the biofilm matrix (compare Fig. 2 C, D, I, J and Fig. 5 G, J), probably, to defend of *P. aeruginosa* antagonistic factors.

By contrast, when the *S. aureus* – *P. aeruginosa* mixed biofilms were treated with any of the broad-spectrum antimicrobials such as ciprofloxacin, gentamicin or amikacin, nearly 10–fold lower concentrations were sufficient to achieve the same reduction in the CFUs number of both bacteria in the biofilm, in comparison with monoculture treatment (Figs 7 and 8, compare violets with reds or blues on panels A and B). This effect was more pronounced for aminoglycosides, which at already 1–2×MBC led to the complete death of both biofilm-embedded and detached cells in dual-species biofilm, while in monocultures 8×MBC was required to reduce the number of CFUs by 3–5 orders of magnitude (Figs 8 and S12) despite of reported synergy of F105 with aminoglycosides ^73^. In the meanwhile, the observed reduction of the *S. aureus* CFUs number was not a consequence of their transition into small colony variants in response to aminoglycosides and cyanide production by *P. aeruginosa* (see Figs S15, S16). Apparently, this effect could be attributed to rhamnolipids synthesis with *P. aeruginosa* which increase *S. aureus* susceptibility to aminoglycosides ^81^.

In contrast to other works on antimicrobial susceptibility of *S. aureus* – *P. aeruginosa* dual species cultures, here we show that *S. aureus* also affects the susceptibility of *P. aeruginosa* biofilm-embedded cells to antimicrobials with the most pronounced effect for aminoglycosides. Moreover, tetracycline and ceftriaxone, while being inefficient against *P. aeruginosa*, at high concentrations significantly reduced the CFUs of this bacterium in the mixed biofilms (Fig. S7). This suggests that mechanisms of interbacterial interaction affecting bacterial susceptibility in biofilms differ from those in liquid cultures and are still to be understood.

Interestingly, under repression of the *S. aureus* biofilm formation by F105, the efficiency of ciprofloxacin and aminoglycosides against *S. aureus* did not change significantly, while the sensitivity of *P. aeruginosa* was restored to the level characteristic for its monoculture biofilm (Figs 7 and 8, compare violets with reds or blues on panels C and D). This effect could be attributed to either the significant reduction of *S. aureus* fraction in the biofilm (see Fig. 9, S13, S14) or the repression of the antagonistic factors production by *S. aureus* due to complex changes in its cell metabolism in the presence of F105 ^73,75^. Nevertheless, the molecular basis of these complex interbacterial interactions that under certain conditions lead to a clear reversal in the antimicrobials susceptibility requires further investigations.

Taken together, our data clearly indicate that efficient treatment of biofilm-associated mixed infections requires antimicrobials which would be active against dominant pathogens. As we have shown for the *S. aureus* and *P. aeruginosa* mixed culture model, in this case the interbacterial antagonism under certain conditions assists antimicrobial treatment. In contrast, treatment by antibiotics with different efficacy against various consortia members leads to the survival of sensitive cells in the matrix formed by the resistant ones.

Finally, we have shown that *S. aureus* and *P. aeruginosa* are able to penetrate into each other’s mature biofilms (see Fig. 1 A and B) and by this intervention significantly affect the susceptibility of the mixed biofilm to antimicrobials (Fig. 10). When *P. aeruginosa* was introduced into *S. aureus* biofilm, all antimicrobials reduced the amount of CFUs of both bacteria in the biofilm by 3 orders of magnitude at 1–2×MBC with more pronounced effect observed for gentamicin. In the reverse experiment, the inoculation of *S. aureus* to the mature *P. aeruginosa* biofilm significantly increased the efficacy of ciprofloxacin against *P. aeruginosa*.

From a broader perspective, we believe that artificial intervention of antagonistic bacteria into already preformed monoculture biofilms could be used to enhance their antimicrobial treatment efficacy. We suggest that this approach has a strong potential of further development towards innovative treatment of biofilm-associated infections such as introduction of the skin residential saprophytic microflora to the biofilms formed by nosocomial pathogens to increase antimicrobial treatment efficacy. While in this work we demonstrated the synergy of interbacterial antagonism with antimicrobials using the well-studied *S. aureus - P. aeruginosa* model system, we believe that many other bacteria of normal body microflora are available to antagonize with nosocomial pathogens and thus can be used for the enhancement of microbial infections treatment by using microbial transplantation.

## Materials and Methods

Derivate of 2(5H)-furanone designed as F105 (3-Chloro-5(S)-[(1R,2S,5R)-2-isopropyl-5-methylcyclohexyloxy]-4-[4-methylphenylsulfonyl]-2(5H)-furanone) was described previously ^83^ and synthesized at the department of Organic Chemistry, A.M. Butlerov Chemical Institute, Kazan Federal University.

### Bacterial strains and growth conditions

*Staphylococcus aureus subsp. aureus* (ATCC 29213) and *Pseudomonas aeruginosa* (ATCC 27853D-5) were used in this assay. The bacterial strains were stored in 10 % (V/V) glycerol stocks at −80 °C and freshly streaked on blood agar plates (BD Diagnostics) followed by their overnight growth at 35 °C before use. Fresh colony material was used to adjust an optical density to 0.5 McFarland (equivalent to 10^8^ cells/mL) in 0.9 % NaCl solution that was used as a working suspension. For the biofilm assay the previously developed BM broth (glucose 5g, peptone 7g, MgSO_4_× 7H_2_O 2.0g and CaCl_2_× 2H_2_O 0.05g in 1.0 liter tap water) ^57,71,84^ where both *S. aureus* and *P. aeruginosa* formed rigid biofilms in 2 days was used. The mannitol salt agar (peptones 10g, meat extract 1g, NaCl 75g, D-mannitol 10g, agar-agar 12g in 1.0 liter tap water, Oxoid) and cetrimide agar (Sigma) were used to distinguish *S. aureus* and *P. aeruginosa*, respectively, in mixed cultures. Bacteria were grown under static conditions at 35°C for 24–72 hours as indicated.

### Construction of *ica*-gfp reporter construction

The 500-bp promoter region of *icaA* gene was amplified from the chromosomal DNA of *S. aureus* USA300 ^85^ by using primers icaA for and icaA rev (table S2), the *gfp* gene was amplified from the pCtuf-gfp plasmid ^27^ by using gfp for and gfp rev (table S2). The PCR products were cloned into the expression vector pRB473 ^86^ digested by HindIII using an isothermal, single-reaction method for assembling multiple overlapping DNA molecules as described previously ^87^ by obtaining a plasmid pRB-ica-gfp. The resulting plasmid was transformed in *S. aureus* ATCC 29213 by electroporation as described previously ^88^.

### Biofilm assays

Biofilm formation was assessed in 24-well polystirol plates (Eppendorf) by staining with crystal violet as described earlier in ^89^ with modifications. Bacteria with an initial density of 3×10^7^ CFU/ml were seeded in 2 ml BM at 37°C and cultivated for 48 h under static conditions. Then the culture liquid was removed and the plates were washed once with phosphate-buffered saline (PBS) pH=7.4 and dried for 20 min. Then, 1 ml of a 0.5% crystal violet solution (Sigma-Aldrich) in 96% ethanol was added per well followed by incubation for 20 min. The unbounded dye was washed off with PBS. The bound dye was eluted in 1 ml of 96% ethanol, and the absorbance at 570 nm was measured on a Tecan Infinite 200 Pro microplate reader (Switzerland). Cell-free wells subjected to all staining manipulations were used as control.

The biofilms were additionally analyzed by confocal laser scanning microscopy (CLSM) on Carl Zeiss LSM 780 confocal microscope. Both mono- and mixed cultures of *S. aureus* and *P. aeruginosa* were grown on cell imaging cover slips (Eppendorf) under static conditions for 48 h in BM broth. Next one-half of the medium was replaced by the fresh one containing antimicrobials at final concentrations as indicated and cultivation was continued for the next 24h. The samples were then stained for 5 min with the SYTO 9 (ThermoFisher Scientific) at final concentration of 0.02 μg/ml (green fluorescence) and propidium iodide (Sigma) at final concentration of 3 μg/ml (red fluorescence) to differentiate between viable and non-viable bacteria. To differentiate between gram-positive and gram-negative bacterial species ViaGram Red^+^ (ThermoFisher Scientific) was used. The microscopic images were obtained with a 1-μm Z-stacks.

### Evaluation of antibacterial activity

The minimum inhibitory concentration (MIC) of antimicrobials was determined by the broth microdilution method in 96-well microtiter plates (Eppendorf) according to the recommendation of the European Committee for Antimicrobial Susceptibility Testing (EUCAST) rules for antimicrobial susceptibility testing ^90^. Briefly, the 10^8^ cells/mL bacterial suspension was subsequently diluted 1:300 with BM broth supplemented with various concentrations of antimicrobials in microwell plates to obtain a 3×10^5^ cells/mL suspension. The concentrations of antimicrobials ranged from 0.25 to 512 mg/L. Besides the usual double dilutions, additional concentrations were included in between. The cultures were next incubated at 35°C for 24 h. The MIC was determined as the lowest concentration of antimicrobials for which no visible bacterial growth could be observed after 24 h incubation.

To determine the MBC of antimicrobials the CFU/mL were further evaluated in the culture liquid from those wells without visible growth. 10 μl of the culture liquid from the wells with no visible growth were inoculated into 3ml of LB broth followed by cultivation for 24h. The MBC was determined as the lowest concentration of compound for which no visible bacterial growth could be observed according to the EUCAST of the European Society of Clinical Microbiology and Infectious Diseases (ESCMID) ^91^.

### Drop plate assay

To evaluate the viability of both detached and planktonic cells, a series of 10-fold dilutions of liquid culture from each well were prepared in 3 technical repeats and dropped by 5 μl onto LB agar plates. CFUs were counted from the two last drops typically containing 5-15 colonies and further averaged. To evaluate the viability of the biofilm-embedded cells, the wells were washed twice with 0.9% NaCl in order to remove the non-adherent cells. The biofilms were also suspended in 0.9% NaCl by scratching the well bottoms with subsequent treatment in an ultrasonic bath for 2 min to facilitate the disintegration of bacterial clumps ^71^. Viable cells were counted by the drop plate method as described above.

### Statistical analysis

Experiments were carried out in six biological repeats with newly prepated cultures and medium in each of them. The fraction of non-viable cells in microscopic images was estimated as the relative fraction of the red cells among all cells in the combined images obtained by overlaying of the green and the red fluorescence microphotographs (10 images per each sample) by using BioFilmAnalyzer software ^92^.The statistical significance of the discrepancy between monoculture and mixed biofilms treatment efficacy was determined using the Kruskal-Wallis statistical test with significance threshold at *p* < 0.05.

## Acknowledgments

This work was supported by the Russian Foundation for Basic Research, grant No 20-04-00247 (AKa), by the Ministry of Education and Science of Russia, assignment No 2019-0460 (MB). This study was partially performed in the framework of the Russian Government Program of Competitive Development of Kazan Federal University.

## Author contributions

ET, MY, DB, FA and ER performed the experiments. AKa and RF conducted the experiments. AKh and AKu synthesized F105. ET, MY, AKa and MB analyzed the results, prepared figures and graphs and wrote the manuscript. All the authors read and approved the final version of the manuscript.

