## Supplementary data for "Bidirectional alterations in antibiotics susceptibility in *Staphylococcus aureus - Pseudomonas aeruginosa* dual-species biofilm"

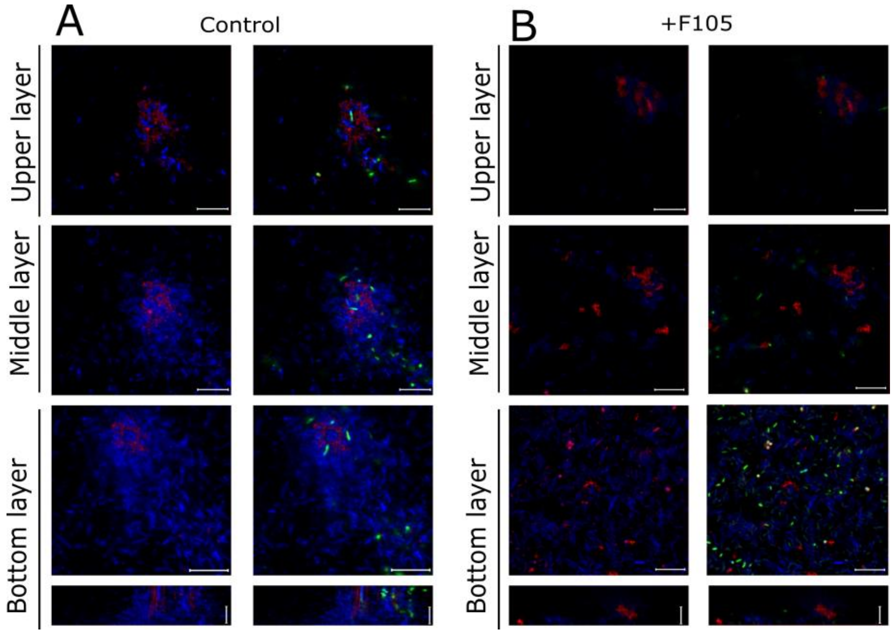


**Figure S1. The distribution and viability of *S. aureus* and *P. aeruginosa* in the mixed biofilm.** Cells were grown without any antimicrobial (A) or in presence of F105 specifically inhibiting the biofilm formation by *S. aureus* cells (B). The 48-h old biofilms were stained by ViaGram Red$^+$ to differentiate *S. aureus* (stained in red), *P. aeruginosa* (stained in blue) and non-viable cells (stained in green) and assessed by CLSM. The CLSM images show a plan view on an upper, middle or bottom biofilm layer and a cross section through the biofilm.

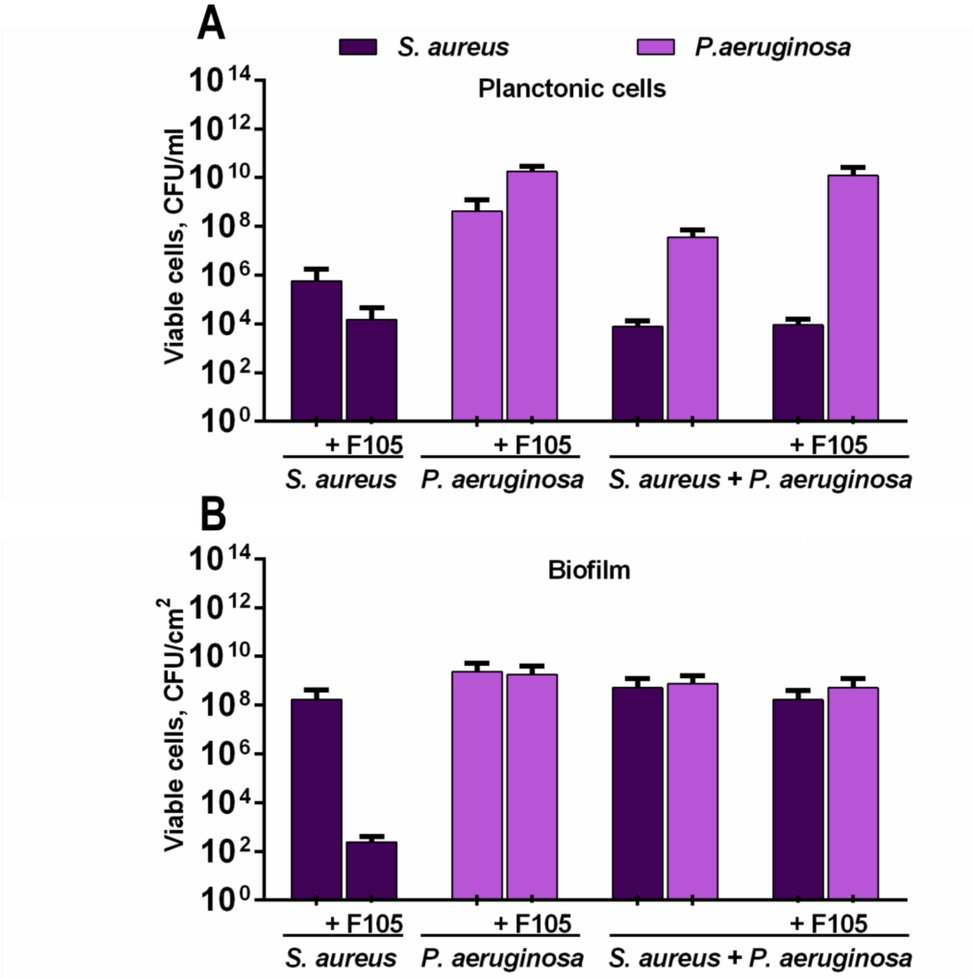


**Figure S2. The number of viable *S. aureus* and *P. aeruginosa* in monomicrobial and mixed cultures (A- planktonic cells, B-biofilm embedded cells) grown in absence or presence of F105.** The 48 hours old biofilms were aseptically washed to remove non-adherent cells and CFUs were counted by drop plate assay. The salt-mannitol agar and cetrimide agar were used to differentiate *S. aureus* and *P. aeruginosa* in mixed biofilms.

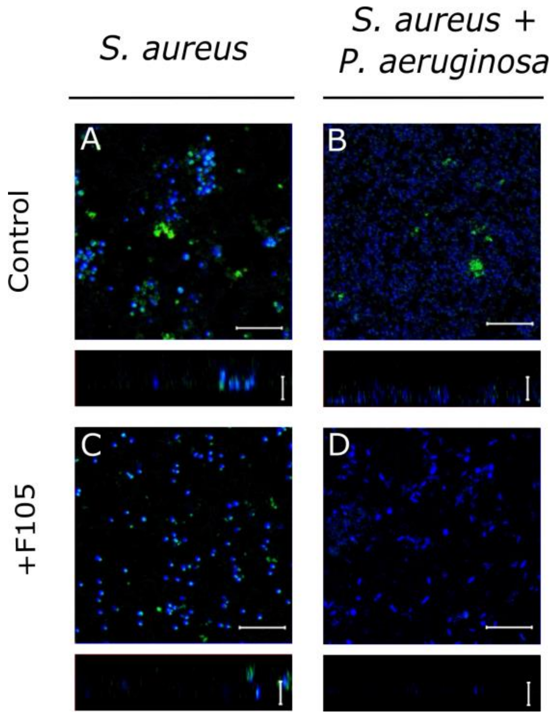


**Figure S3. Evaluation of the *ica*-GFP repression in *S. aureus* cells in presence of F105 in mono- and mixed biofilms.** Cells were grown in absence (A, B) or in presence (C, D) of 2(5*H*)-furanone derivative F105 specifically inhibiting the biofilm formation by *S. aureus* cells. The 48-h old biofilms were assessed by CLSM. The scale bars indicate 10 µm.

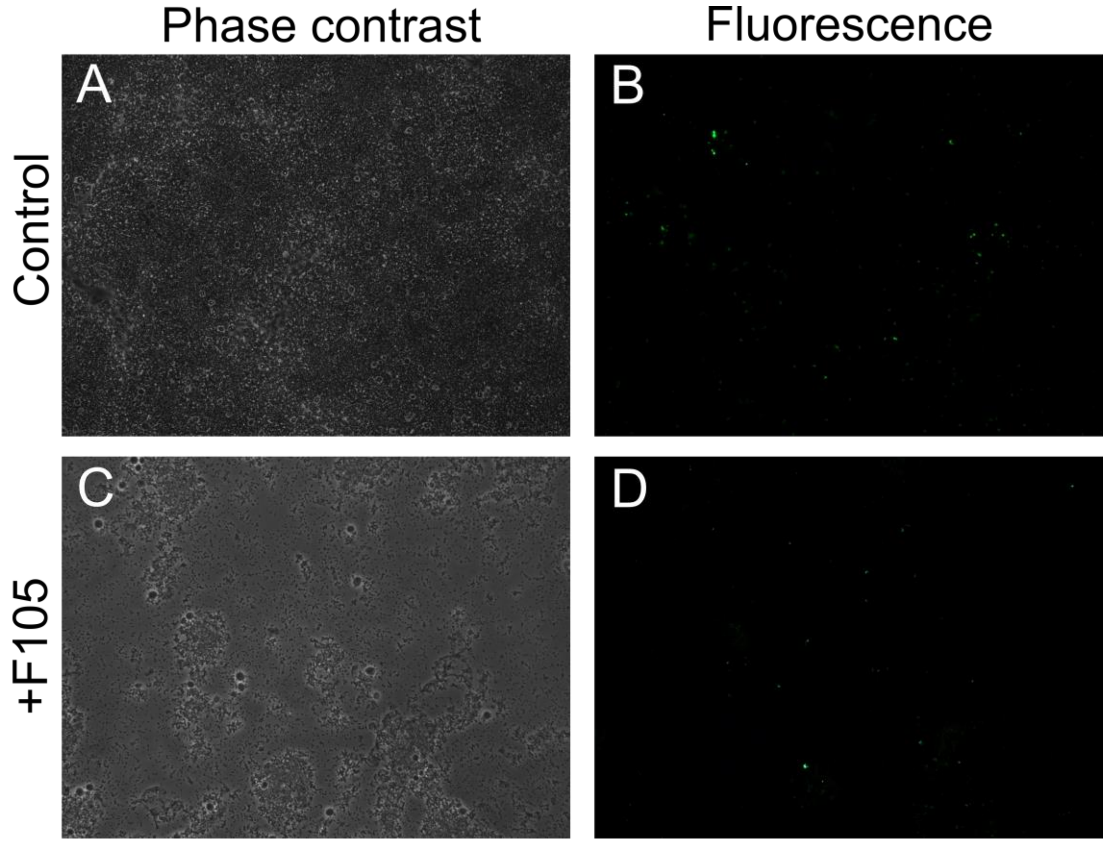


**Figure S4. Evaluation of the constitutive expression of GFP in *S.aureus* pC-tuf-gfp cells in presence of F105 in biofilm.** Cells were grown in absence (A, B) or in presence (C, D) of 2(5*H*)-furanone derivative F105 specifically inhibiting the biofilm formation by *S. aureus* cells. The 48-h old biofilms were assessed by fluorescence microscopy.

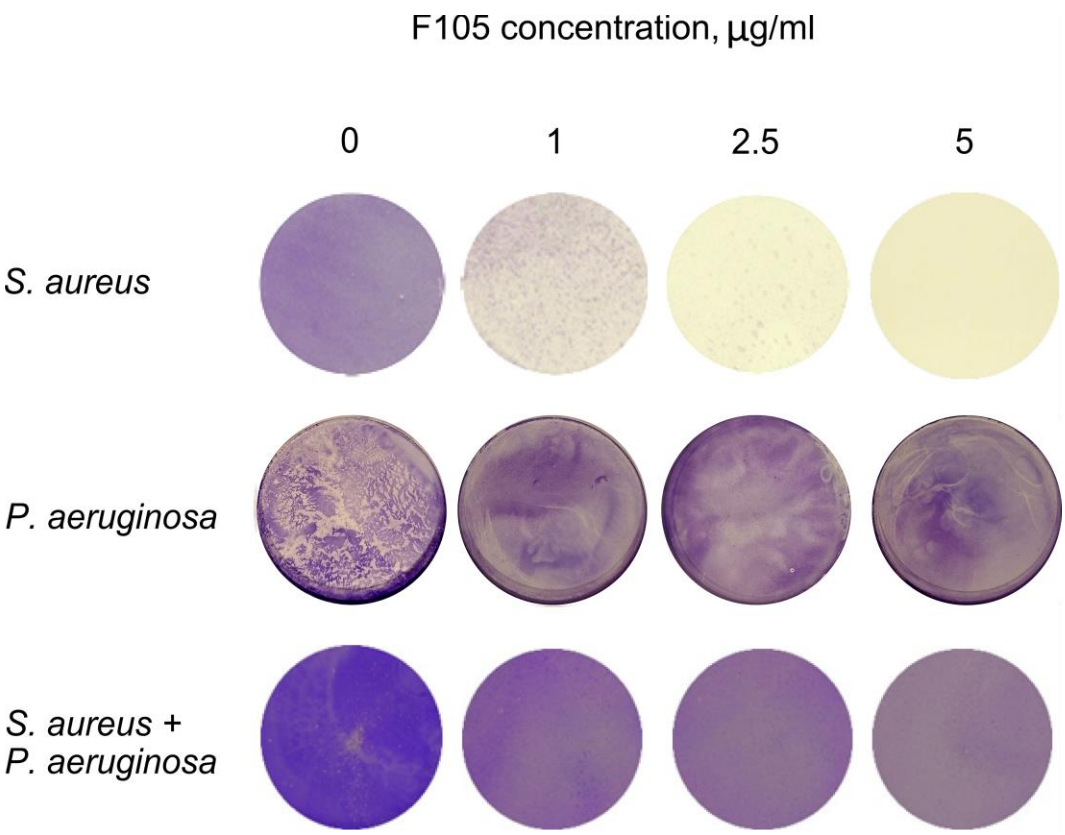


**Figure S5. The effect of 2(5H)-furanone derivative (F105) on the formation of monomicrobial biofilms of *S. aureus* and *P. aeruginosa* and mixed one.** The biofilms were assessed with crystal-violet staining, the bottoms of well are shown.

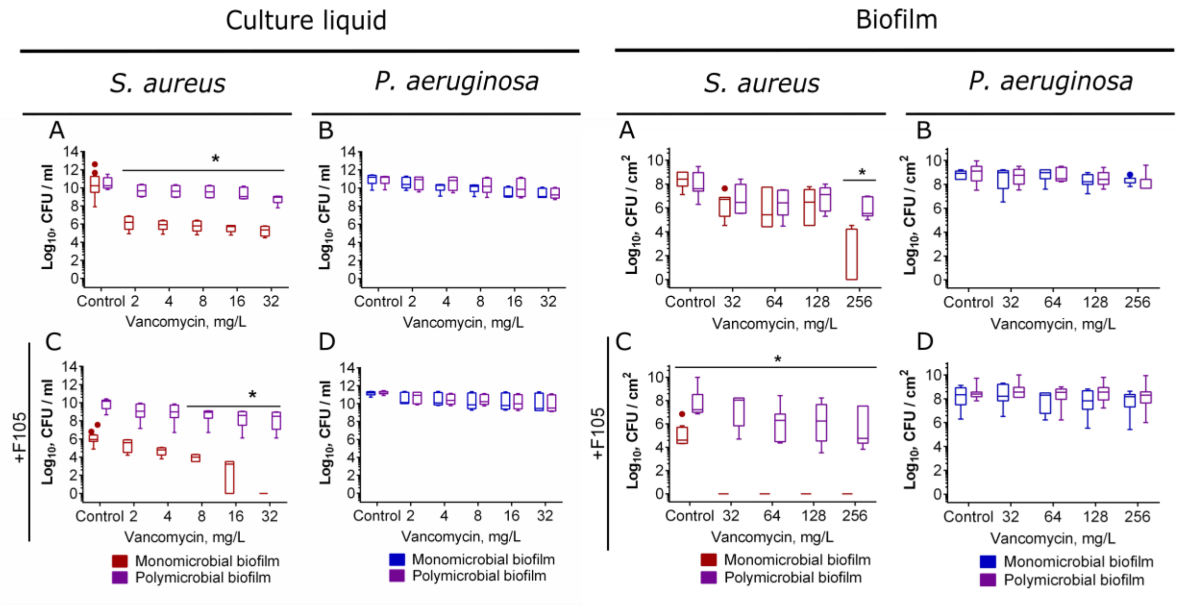


**Figure S6. The effect of vancomycin on viability of *S. aureus* and *P.aeruginosa* detached cells and biofilm embedded cells into their mono- and polymicrobial biofilms.** Antimicrobials were added to 48 hours-old biofilms grown in absence (A-B) or presence (C-D) of F105 to inhibit the biofilm formation by *S. aureus*. After 24 h incubation, the biofilms were washed twice with sterile 0.9% NaCl. The adherent cells were scratched, resuspended and their viability was analyzed by using drop plate assay. Asterisk shows significant difference between CFUs number in monomicrobial and mixed biofilms.

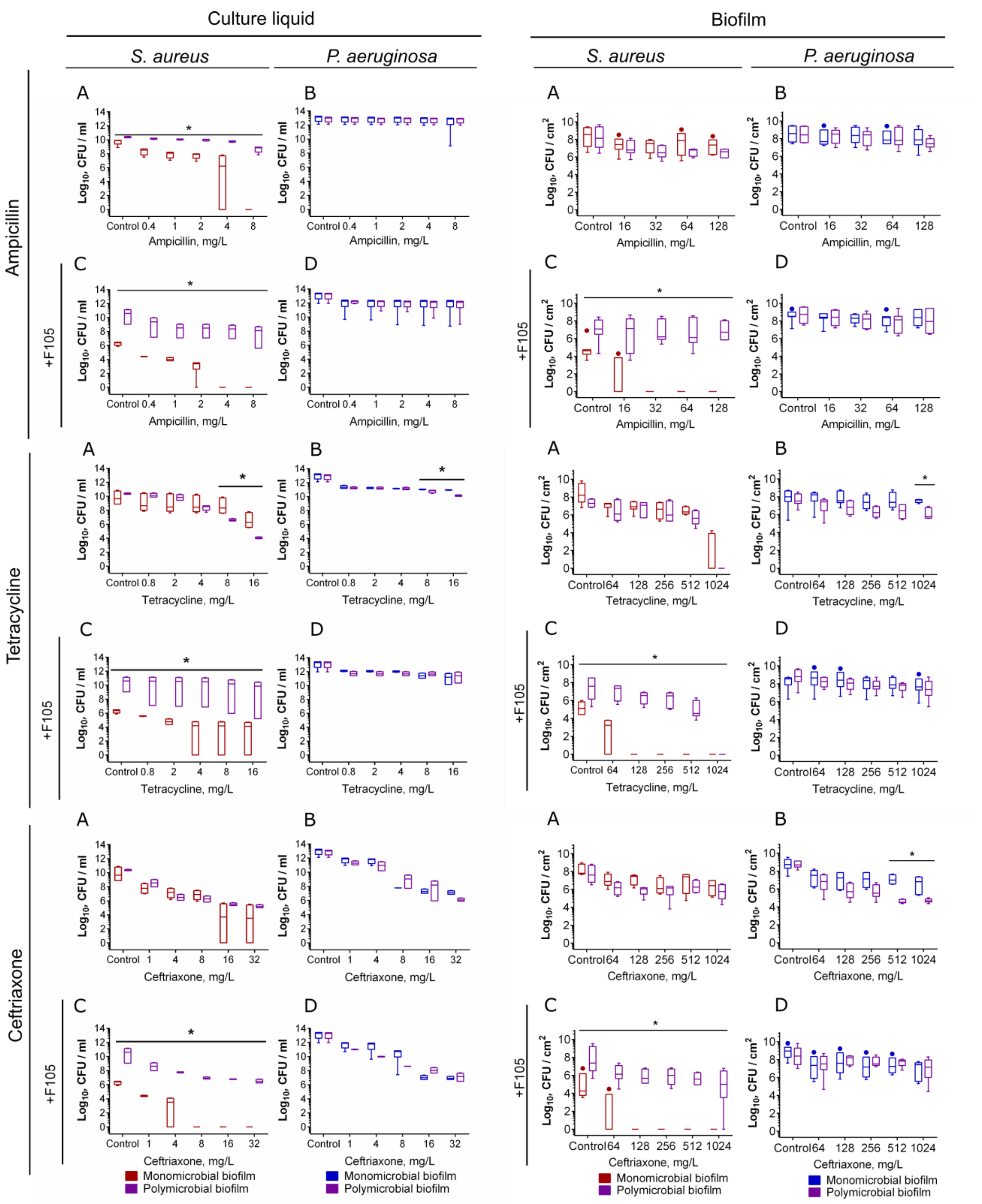


**Figure S7. The effect of ampicillin, tetracycline and ceftriaxone on viability of *S. aureus* and *P.aeruginosa* detached planktonic cells and biofilm embedded cells into their mono- and polymicrobial biofilms.** Antimicrobials were added to 48 hours-old biofilms grown in absence (A-B) or presence (C-D) of F105 to inhibit the biofilm formation by *S. aureus*. After 24 h incubation, the biofilms were washed twice with sterile 0.9% NaCl. The adherent cells were scratched, resuspended and their viability was analyzed by using drop plate assay. Asterisk shows significant difference between CFUs number in monomicrobial and mixed biofilms.

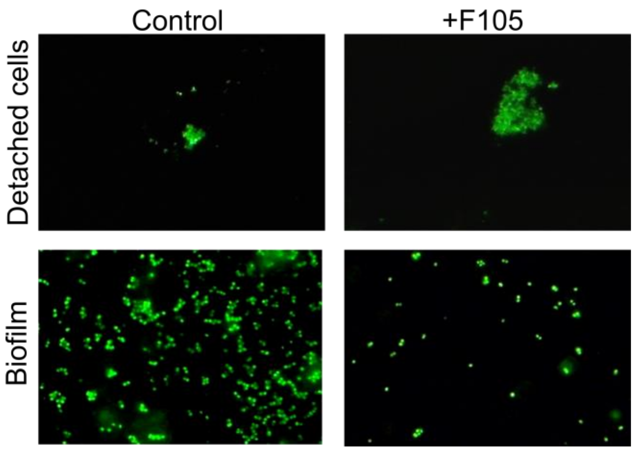


**Figure S8. The effect of F105 on viability of *S. aureus* detached planktonic cells clumps and biofilm embedded cells into their monomicrobial biofilm.** Cells were grown in absence (A, B) or in presence (C, D) of 2(5*H*)-furanone derivative F105 specifically inhibiting the biofilm formation by *S. aureus* cells. The 48-h old biofilms were assessed by fluorescence microscopy.

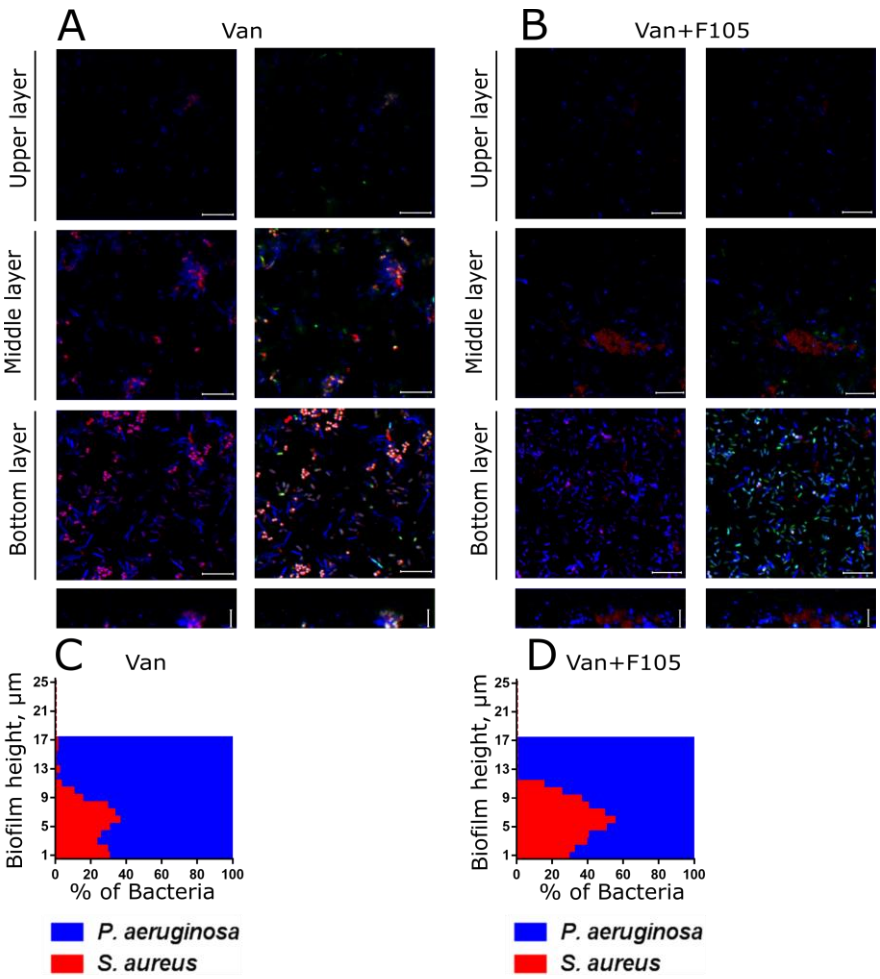


**Figure S9. The effect of vancomycin on *S. aureus* and *P. aeruginosa* viability and distribution in mixed biofilms grown in absence (A, C) or in presence of F105 specifically inhibiting the biofilm formation by *S. aureus* cells (B, D).** Vancomycin (256 µg/mL corresponding to 8×MBC for *S. aureus*) was added to 48 hours-old biofilms. After 24 h incubation, the biofilms were stained by ViaGram Red+ to differentiate *S. aureus* (stained in red), *P. aeruginosa* (stained in blue) and non-viable cells (stained in green) and assessed by CLSM. The images show a plan view on an upper, middle or bottom biofilm layer and a cross section through the biofilm. The scale bars indicate 10 µm.

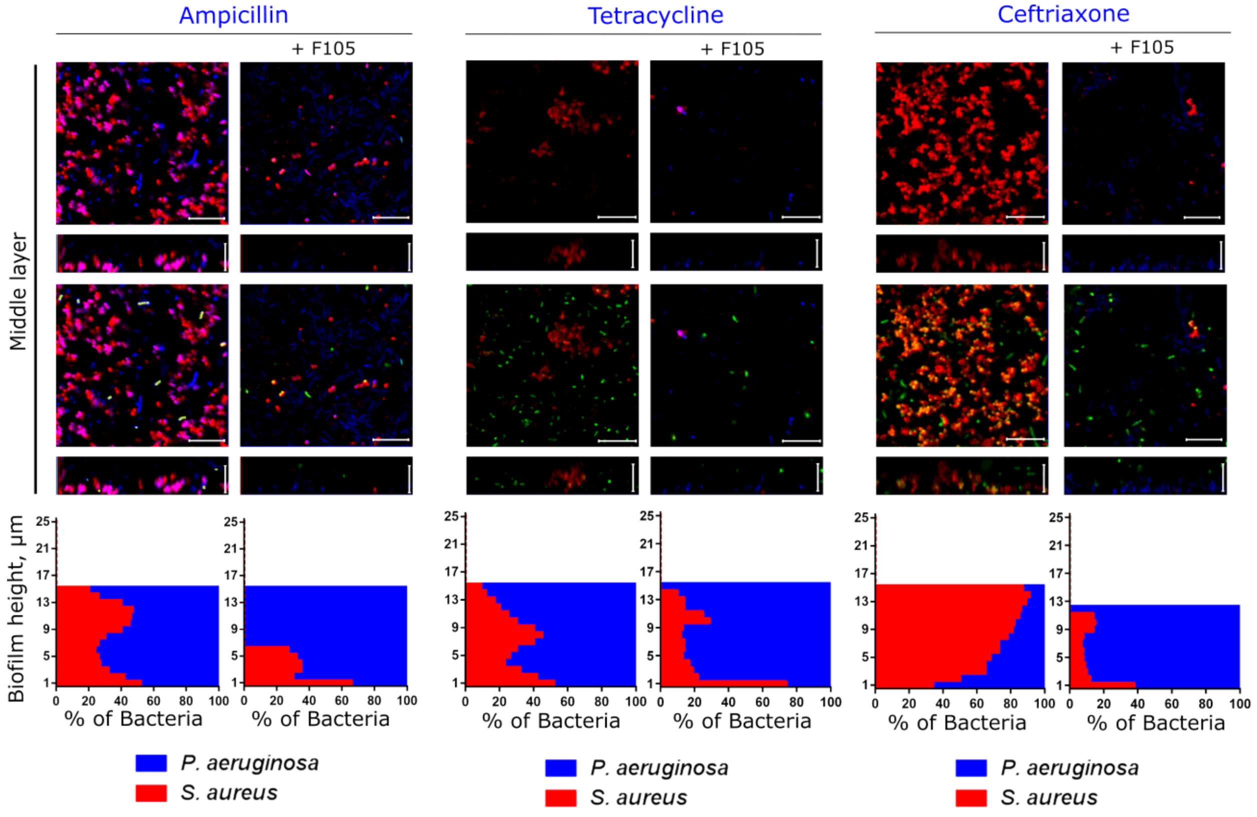


**Figure S10. The effect of tetracycline, ampicillin and ceftriaxone on viability and distribution of *S. aureus* and *P. aeruginosa* in mixed biofilms grown normally or in presence of F105 specifically inhibiting the biofilm formation by *S. aureus* cells.** Antimicrobials (at concentrations corresponding to their 8×MBC for S. aureus, see Table 1 for values) were added to 48 hours-old biofilms. After 24 h incubation, the biofilms were stained by ViaGram Red+ to differentiate *S. aureus* (stained in red), *P. aeruginosa* (stained in blue) and non-viable cells (stained in green) and assessed by CLSM. The images show a plan view on a middle biofilm layer (indicated by arrows) and a cross section through the biofilm. The scale bars indicate 10 µm. The distributions of *S. aureus* and *P. aeruginosa* in the biofilm layers assessed from the CLSM image layers by using in-house developed BioFilmAnalyser software are expressed as their relative fractions.

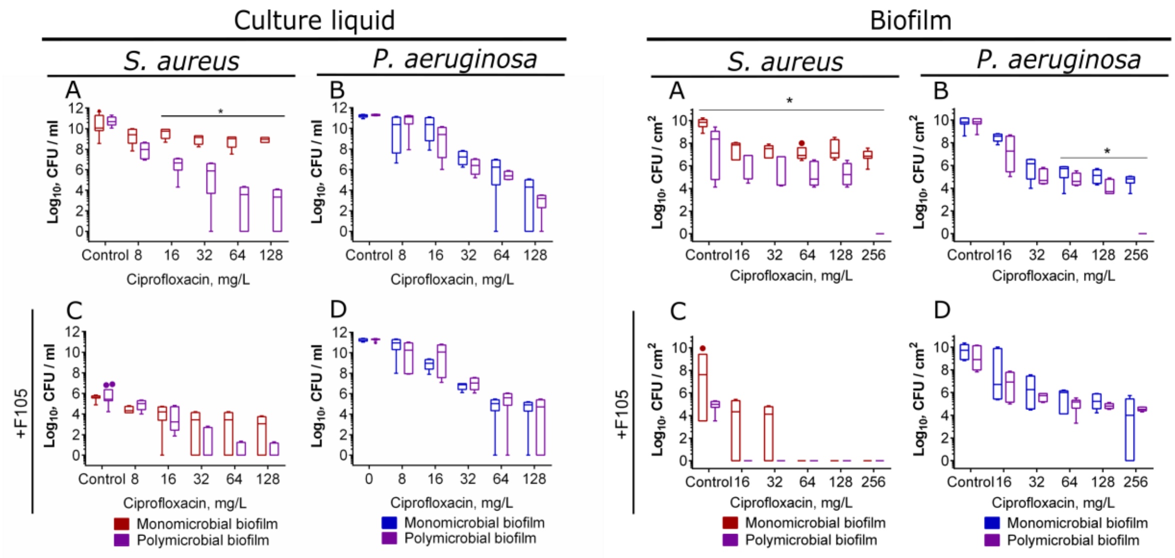


**Figure S11. The effect of ciprofloxacin on viability of *S. aureus* and *P.aeruginosa* detached planktonic cells and biofilm embedded cells into their mono- and polymicrobial biofilms.** Antimicrobials were added to 48 hours-old biofilms grown in absence (A-B) or presence (C-D) of F105 to inhibit the biofilm formation by *S. aureus*. After 24 h incubation, the biofilms were washed twice with sterile 0.9% NaCl. The adherent cells were scratched, resuspended and their viability was analyzed by using drop plate assay. Asterisk shows significant difference between CFUs number in monomicrobial and mixed biofilms.

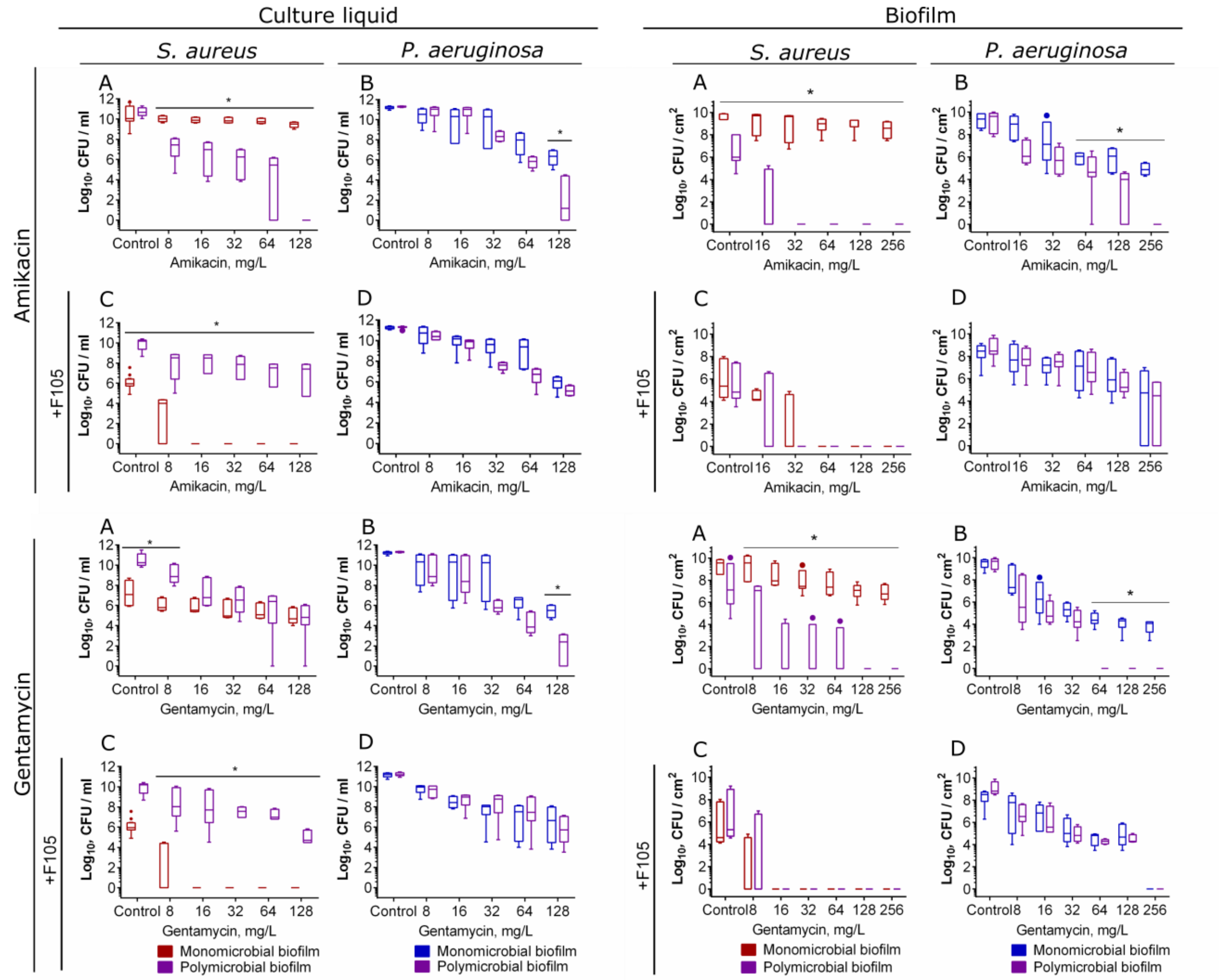


**Figure S12. The effect of aminoglycosides on viability of *S. aureus* and *P.aeruginosa* detached planktonic cells and biofilm embedded cells into their mono- and polymicrobial biofilms.** Antimicrobials were added to 48 hours-old biofilms grown in absence (A-B) or presence (C-D) of F105 to inhibit the biofilm formation by *S. aureus*. After 24 h incubation, the biofilms were washed twice with sterile 0.9% NaCl. The adherent cells were scratched, resuspended and their viability was analyzed by using drop plate assay. Asterisk shows significant difference between CFUs number in monomicrobial and mixed biofilms.

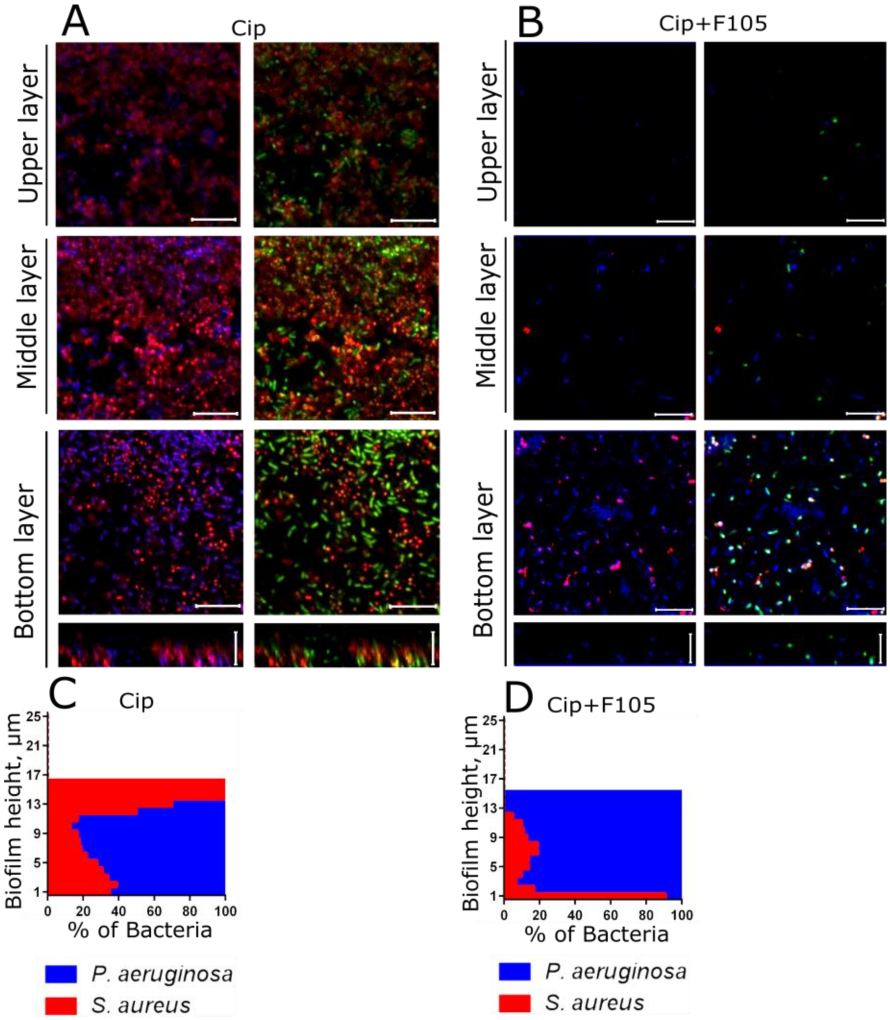


**Figure S13. The effect of ciprofloxacin on viability and distribution of *S. aureus* and *P. aeruginosa* in mixed biofilms grown normally (A) or in presence of F105 specifically inhibiting the biofilm formation by *S. aureus* cells (B).** Ciprofloxacin (512 µg/mL corresponding to 8×MBC for S. aureus) was added to 48 hours-old biofilms. After 24 h incubation, the biofilms were stained by ViaGram Red+ to differentiate *S. aureus* (stained in red), *P. aeruginosa* (stained in blue) and non-viable cells (stained in green) and assessed by CLSM. The images show a plan view on an upper, middle or bottom biofilm layer (indicated by arrows) and a cross section through the biofilm. The scale bars indicate 10 µm.

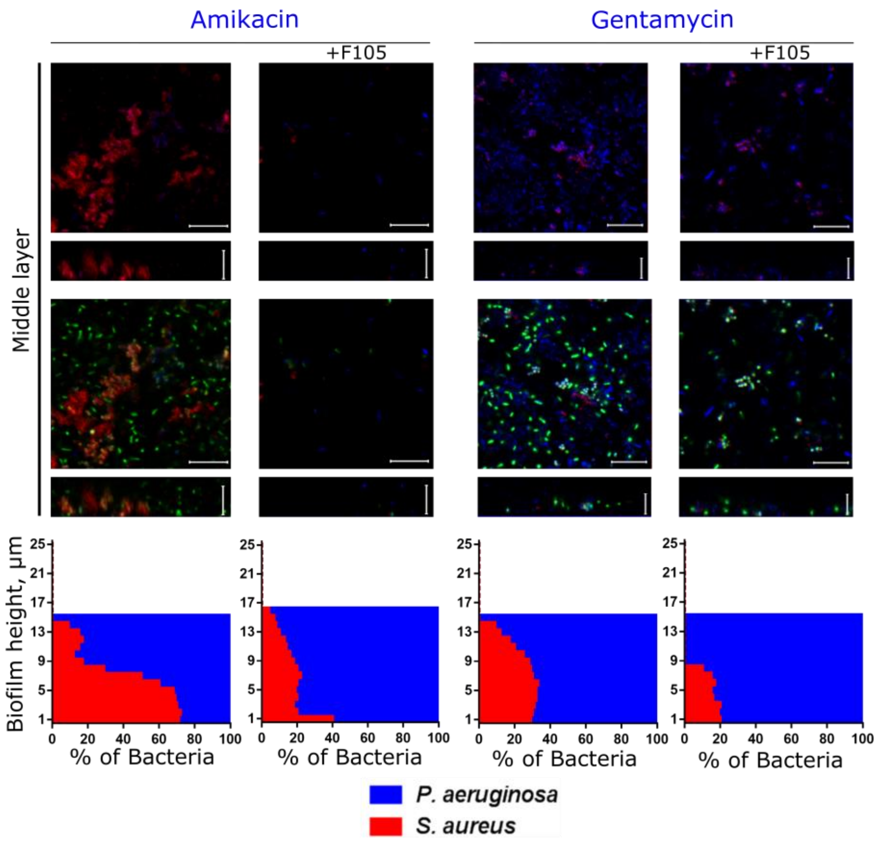


**Figure S14. The effect of amikacin and gentamycin on viability and distribution of *S. aureus* and *P. aeruginosa* in mixed biofilms grown normally or in presence of F105 specifically inhibiting the biofilm formation by *S. aureus* cells.** Antimicrobials (512 µg/mL, corresponding to 8×MBC for both *S. aureus* and *P. aeruginosa*) were added to 48 hours-old biofilms. After 24 h incubation, the biofilms were stained by ViaGram Red+ to differentiate *S. aureus* (stained in red) and P.aeruginosa (stained in blue) and biofilms were assessed by CLSM. The images show a plan view on a middle biofilm layer (indicated by arrows) and a cross section through the biofilm. The scale bars indicate 10 µm. The distribution of *S. aureus* and *P. aeruginosa* in the biofilm layers assessed from the CLSM image layers by using in-house developed BioFilmAnalyser software are expressed as their relative fractions.

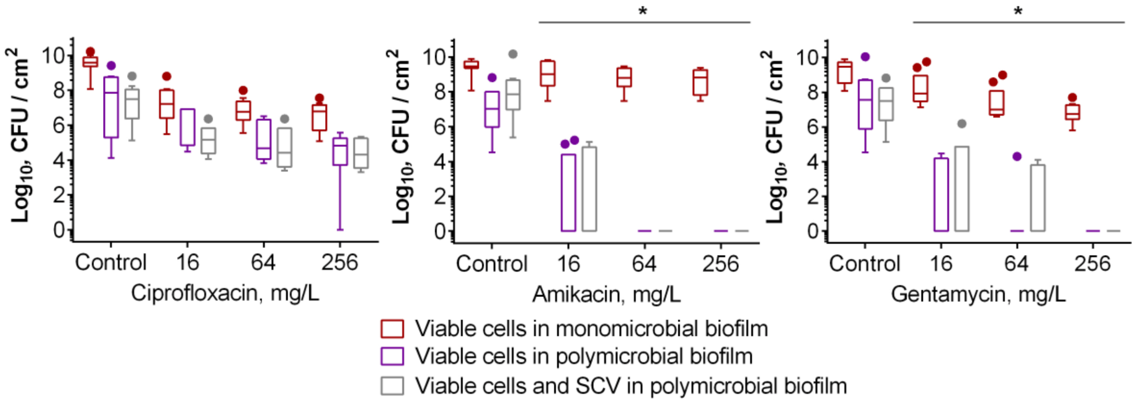


**Figure S15. The evaluation of frequency of *S. aureus* transition to Small Colony Variants in mixed biofilms when treated with broad-spectrum antibiotics.** Antimicrobials were added to 48 hours-old *S. aureus* and *P. aeruginosa* mixed biofilms. After 24 h incubation, the biofilms were washed twice with sterile 0.9% NaCl. The adherent cells were scratched, resuspended and CFUs were counted. To count *S. aureus* SCV, the cell suspensions were seeded onto LB plates with colistin (32 µg/mL) and grown for 5 days (grey columns).

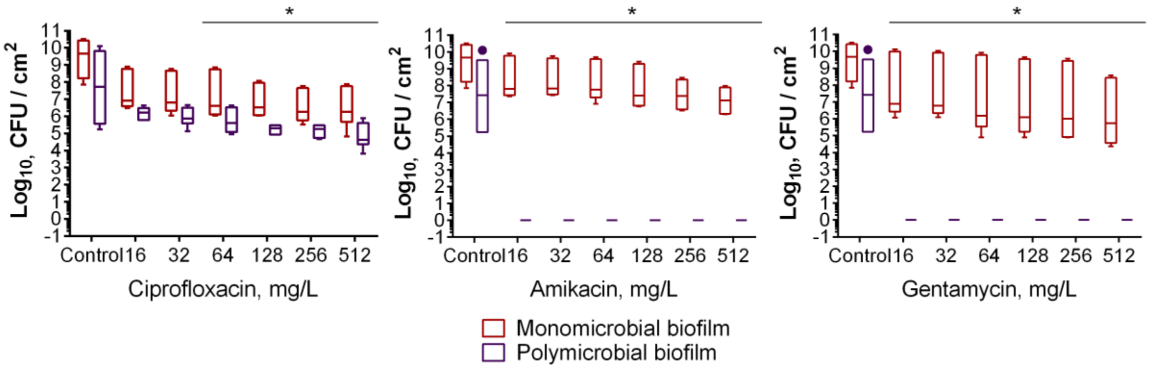


**Figure S16. The effect of broad-spectrum antibiotics on viability of cyanide-resistant *S. aureus* 0349 pCXcydAB$_{sa}$ strain in polymicrobial biofilms.** Antimicrobials were added to 48 hours-old biofilms. After 24 h incubation, the biofilms were washed twice with sterile 0.9% NaCl. The adherent cells were scratched, resuspended and CFUs were counted.

**Table S1. ECOFF, MIC and MBC values in μg/mL of various antibiotics against *S. aureus* and *P. aeruginosa*.**

| | *S. aureus* | | | *P. aeruginosa* | | |
|---|---|---|---|---|---|---|
| | ECOFF | MIC | MBC | ECOFF | MIC | MBC |
| F105 | ND | 2.5 | 5 | ND | ND | ND |
| Van | 2.0 | 4 | 32 | ND | ND | ND |
| Tet | 1.0 | 0.25 | 128 | ND | 16 | ND |
| Cef | 8.0 | 8 | 128 | ND | 32 | ND |
| Amp | ND | 0.5 | 16 | ND | ND | ND |
| Ami | 8.0 | 2 | 64 | 16 | 1 | 64 |
| Gen | 2.0 | 4 | 32 | 8.0 | 8 | 64 |
| Cip | 1.0 | 1 | 64 | 0.5 | 4 | 64 |

MIC and MBC were assessed by the broth microdilution.
*ND – not determined

**Table S2. Primers for *ica*-gfp reporter construction.**

| Primer | Sequense |
|---|---|
| *icaA* for | 5'- CGT TTT TTA TTG GTG AGA ATC CAA GCT TGT CCG TAA ATA TTT CCA GAA AAT TC - 3' |
| *icaA* rev | 5'-TAG CGC TCA TTT TCT TTA CCT ACC TTT CG - 3' |
| gfp for | 5'- GGT AAA GAA AAT GAG CGC TAG CAA AGG AG - 3' |
| gfp rev | 5'- AGT CGA CCT GCA GGC ATG CAA GCT TTG TAT AGT TCA TCC ATG CCA TG - 3' |
